# PIP2 Binding at Allosteric Site Blocks Activation in Human Rod CNG Channels

**DOI:** 10.64898/2025.12.19.695557

**Authors:** Taehyun Park, Crina M. Nimigean

## Abstract

Phosphatidylinositol-4,5-bisphosphate (PIP2) is a ubiquitous signaling lipid that regulates multiple ion channels. In human cyclic nucleotide-gated (CNG) channels, including the rod photoreceptor channel, PIP2 has been reported to exert inhibitory effects, but the underlying mechanism has remained unclear. Because this inhibition lowers the apparent cGMP sensitivity of rod CNG channels, it can play a key role in controlling the light sensitivity and dynamic range of rod photoreceptors. Here we report how PIP2 modulates the function of human CNGA1 channels, the major subunit of human rod photoreceptor CNG channels. Ensemble ion flux assays with liposome-reconstituted purified CNGA1 channels demonstrated robust inhibition by PIP2 via a reduction in apparent cGMP sensitivity, and single-channel recordings revealed PIP2 reduces the channel’s open probability without altering unitary conductance. To uncover the structural basis, we determined cryo-EM structures of CNGA1 in lipid nanodiscs under multiple ligand conditions. In PIP2-free conditions, closed, intermediate, and open conformations were observed, whereas in the presence of PIP2, the open state was absent. Density consistent with bound PIP2 was detected at inter-protomer grooves between the voltage-sensing and pore domains indicating that PIP2 binding stabilizes non-conductive conformations by sterically preventing C-linker elevation and outward movement of helix S6, conformational changes needed for pore dilation. Collectively, our results establish a structural mechanism for PIP2-mediated inhibition of rod CNG channels, define a mechanistic framework for phosphoinositide control of ligand-gated channels across the CNG superfamily, and provide an inhibitory allosteric binding site for future drug targeting in this channel family.

## Introduction

Rod cyclic nucleotide–gated (CNG) channels are non-selective cation channels that allow the passage of Na⁺, K⁺, and Ca²⁺ ions^1^. Although they contain a voltage sensing domain (VSD), their gating is only weakly voltage-dependent and is instead primarily governed by cyclic guanosine monophosphate (cGMP)^2,3^. In rod photoreceptors, CNG channels represent the final transducers of the phototransduction cascade^4^. In darkness, intracellular cGMP is maintained at micromolar levels by the balanced activity of guanylyl cyclases (GC) and phosphodiesterase-6 (PDE6) ^5^. A fraction of CNG channels therefore remain open, producing the “dark current” carried mainly by Na⁺ and Ca²⁺^4,6^. This current sets the resting membrane potential and sustains tonic Ca²⁺ influx at the synaptic terminal. Upon illumination, activation of the rhodopsin–transducin cascade stimulates PDE6, lowering cytosolic cGMP^7^. The resulting closure of CNG channels hyperpolarizes the photoreceptor cell membrane, which in turn reduces Ca²⁺ entry through CaV1.4 L-type Ca²⁺ channels at the ribbon synapse and diminishes glutamate release^8,9^. In this way, changes in CNG channel gating in the outer segment are faithfully transmitted to the synaptic terminal through shifts in membrane potential, initiating signal transmission to bipolar and horizontal cells. Thus, modulation of rod CNG channel activity has profound physiological consequences, shaping the sensitivity, dynamic range, and recovery of rod-mediated vision and preventing response saturation under fluctuating light conditions^10^.

Mammalian rod CNG channels assemble as heterotetramers composed of three CNGA1 subunits and one CNGB1 subunit^11,12^. The obligatory CNGB1 subunit in native channels confers gating, permeation, and pharmacological properties that distinguish heterotetramers from CNGA1 homotetramers, which can form stable, functional but non-native channels in heterologous expression systems^13^. Structural studies using cryo-electron (cryo-EM) microscopy have shown that CNG channels adopt a non–domain-swapped architecture, in contrast to many other tetrameric ion channels^6,14–16^. In rod and cone CNG channels, as well as in evolutionary relatives such as the *C. elegans* TAX-4/TAX-2 and the bacterial channel SthK, cyclic nucleotides (cGMP or cAMP) bind to the CNBD, inducing upward displacement of the cytosolic gating ring, including the C-linker, and the resulting motion is transmitted to the transmembrane gate, where dilation and rotation of the S6 helices open the pore^17–19^. This CNBD–C-linker–to-gate coupling mechanism is highly conserved across the CNG/HCN channel superfamily and underlies the conversion of ligand binding into ion conduction.

Rod CNG channels are subject to multiple layers of regulation that tune their responses to the visual environment. One central feedback pathway involves the Ca²⁺-binding protein calmodulin (CaM), which associates with two distinct sites on the CNGB1 subunit^20,21^ and, upon Ca²⁺ binding, lowers the apparent affinity of the channel for cGMP^20,22^. This desensitization accelerates photoresponse recovery and contributes to light adaptation^22^. Another major axis of modulation is provided by the lipid environment. Among membrane phosphoinositides, phosphatidylinositol 4,5-bisphosphate (PIP2) has been reported to inhibit rod CNG channels, reducing macroscopic current amplitudes and shifting the activation curve toward higher cGMP concentrations^23^. PIP2 is present at low levels in the rod outer segment plasma membrane and undergoes light-dependent turnover rather than remaining static^24–26^. However, the specific molecular interfaces through which PIP2 acts in rods have not been fully resolved. In cone CNG channels, both PIP2 and phosphatidylinositol 3,4,5-trisphosphate (PIP3) exert similar inhibitory effects^27^, whereas in olfactory CNG channels, PIP3 suppresses opening^28^. By contrast, in HCN channels, PIP2 facilitates opening by shifting the voltage dependence of activation toward depolarized potentials^29,30^. Mechanistic insights revealing specific PIP2 binding interactions have recently come from studies of SthK, a bacterial homolog^19,31^. Together, these findings highlight phosphoinositides as versatile regulators with divergent effects across related channels, although the molecular basis by which phosphoinositides regulate human CNG channels remains poorly understood.

Here, to understand how PIP2 interacts with rod CNG channels and alters the cGMP-dependent gating cycle, we investigated the mechanism of PIP2 modulation in CNGA1 channels. To isolate lipid effects, we purified human CNGA1 channel complexes, reconstituted them into proteoliposomes and lipid nanodiscs, and performed functional and structural analyses. Flux assays revealed concentration-dependent inhibition of purified CNGA1 channels by PIP2, while single-channel recordings showed that PIP2 inhibits by decreasing CNGA1 open probability. Cryo-EM structures of nanodisc-embedded CNGA1 channels identified the binding mode of PIP2, captured an intermediate conformation between open and closed states, and revealed how PIP2 binding prevents the transition to an open state. Our findings provide the first structural elucidation of phosphoinositide modulation in human CNG channels and establish a framework for understanding lipid regulation across the broader CNG/HCN superfamily. The identification of a novel allosteric inhibition site for this channel family is essential toward future drug discovery exploits.

## Results

### PIP2 lowers the apparent cGMP sensitivity of CNGA1 channels

To rigorously test the mechanism by which PIP2 inhibits CNGA1 activity, we purified human CNGA1 protein and reconstituted it into proteoliposomes where we can precisely control the amount of PIP2 incorporated (Supplementary Figure 1), thereby eliminating confounding cellular factors that complicated earlier reports of PIP2 inhibition in rod CNG channels^23^. For flux assays, proteoliposomes were loaded with the fluorophore 8-aminonaphthalene-1,3,6-trisulfonic acid (ANTS), which undergoes fluorescence quenching upon Tl⁺ influx through open channels (Figure 1A)^32^. The lipid composition of the rod outer segment (ROS) plasma membrane has been reported as 65% DOPC, 11% POPE, and 24% POPS^33^. Using this mixture as a base, we also prepared proteoliposomes in which DOPC was partially substituted with 0.2%, 1.0%, or 2.5% PIP2. In CNGA1 proteo-liposomes without PIP2, cGMP elicited rapid fluorescence decay, reflecting robust Tl⁺ influx (Figure 1B). A cGMP dose-response curve yielded an EC50 for channel activation with cGMP (∼140 μM) that is higher but on the same order of magnitude as values reported for CNGA1 channels in cells (EC50 ≈ 20–60 µM)^2,34^, indicating that the purified CNGA1 channel was intact and functional (Figure 1C). Increasing the fraction of PIP2 progressively slowed the decay rate, indicating inhibition of channel opening in a PIP2 concentration-dependent manner (Figure 1B). Quantitative analysis revealed that the apparent EC^50^ for cGMP activation shifted from ∼140 µM in no PIP2 to ∼150 µM in 0.2%, ∼190 µM in 1%, and ∼220 µM in 2.5% PIP2 (Figure 1C). These results demonstrate that PIP2 directly inhibits CNGA1 channels, lowering channel activity at physiological cGMP concentrations.

**Figure 1.**
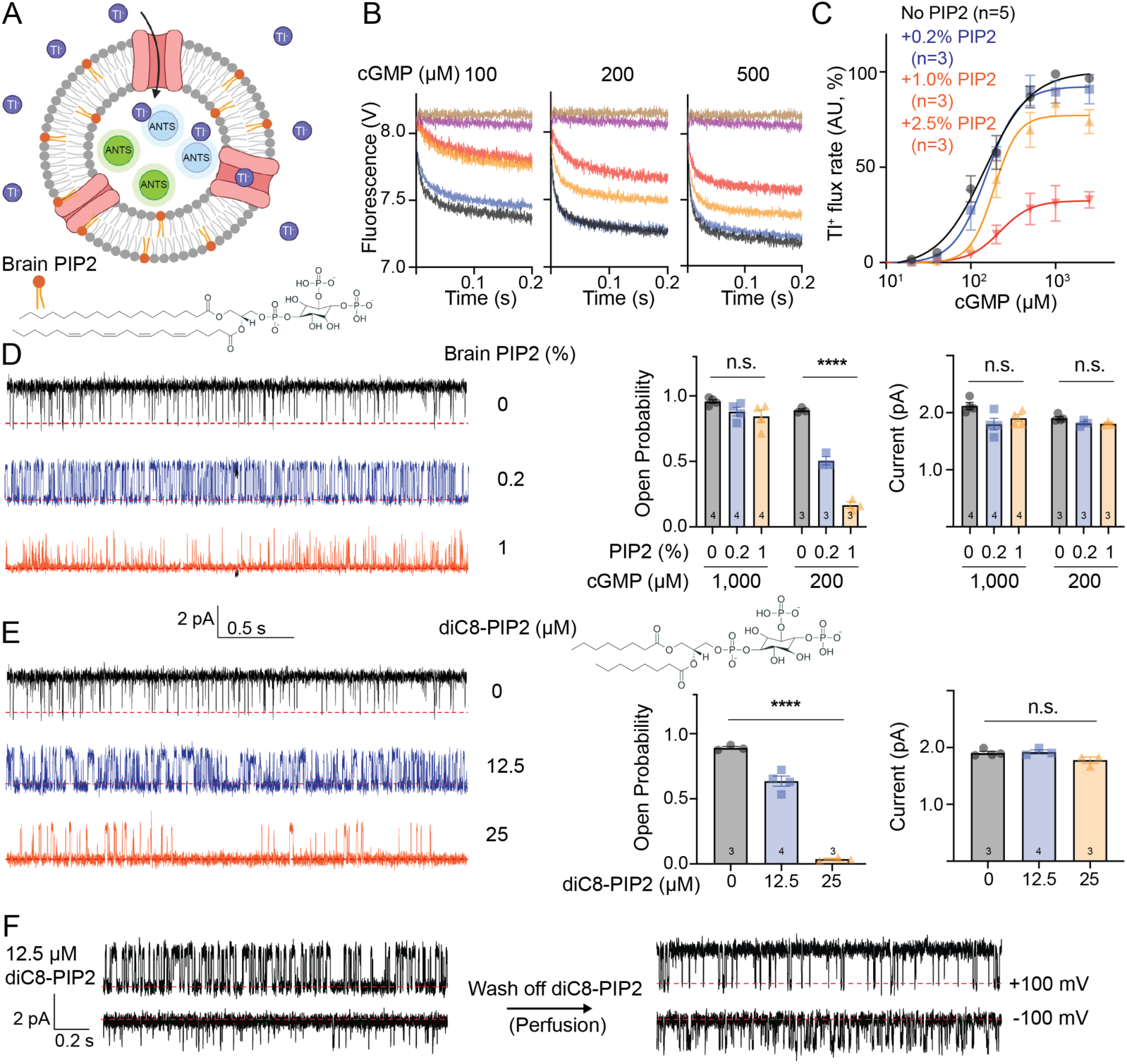
PIP2 inhibits CNGA1 activity in a cGMP-dependent manner. (A) Diagram of the Tl^+^ influx assay using CNGA1 proteoliposomes encapsulating the fluorophore ANTS (green). When cGMP is added, CNGA1 channels open allowing Tl^+^ (purple) influx. Tl^+^ binding to ANTS quenches its fluorescence (blue). Brain PIP2 (orange and chemical structure) was added to liposomes as indicated. (B) Representative fluorescence quenching traces from CNGA1 liposomes with 0 (black), 0.2 (blue), 1.0 (orange), and 2.5 (red) % brain PIP2 at the indicated cGMP concentrations. Brown, buffer only; purple, quencher only without cGMP. (C) Dose–response relationship of Tl⁺ influx rates as a function of cGMP concentration under varying PIP2 conditions. Curves show fits to the Hill equation. The fitted EC50 values in μM for 0, 0.2%, 1.0%, and 2.5% PIP2 were 146.3 ± 18.3, 152.7 ± 18.1, 187.4 ± 22.2, and 220.5 ± 86.6 (mean ± SE), respectively. (D) Left: Representative planar lipid bilayer single-channel recordings of CNGA1 at +100 mV, 200 μM cGMP, with increasing mole fractions of brain PIP2, indicated. Right: Open probabilities and single-channel current amplitudes at indicated cGMP and brain PIP2 from data such as on the left (+100 mV). (E) Left: Representative planar lipid bilayer single-channel recordings of CNGA1 at +100 mV, 200 μM cGMP with soluble diC8-PIP2 in trans chamber. Right: Open probabilities and single-channel current amplitudes at indicated diC8-PIP2 concentrations from data such as on the left (+100 mV). (F) Reversibility of diC8-PIP2 inhibition. Representative single-channel traces for CNGA1 in 200 μM cGMP before and after perfusion of 12.5 μM diC8-PIP2. Closed-channel baseline is indicated by a red dotted line. Bars show mean ± SEM; each dot represents one independent bilayer recording. The numbers on the bars indicate the independent bilayers for each condition. Statistics: conditions were compared by one-way ANOVA followed by Tukey’s multiple-comparison test; ****p < 0.0001, n.s. = not significant. The p values of Po in (D) and (E) were 2.35 × 10^-6^, 5.47×10^-7^, respectively. Recording buffer was 10 mM HEPES and 100 mM KCl (pH 7.4).

To gain more insight into how PIP2 inhibits CNGA1 channels, we performed planar bilayer single-channel recordings in the presence or absence of brain PIP2. This approach enabled direct visualization of gating transitions from single molecules in real time, with orientation controlled by adding ligand to only one chamber of the bilayer (Supplementary Figure 2A). Under control lipid conditions, without PIP2, CNGA1 displayed robust activity in saturating cGMP concentrations (1 mM cGMP), exhibiting an open probability (Pₒ) of ∼1.0 and a unitary current amplitude of ∼2 pA at +100 mV, corresponding to a conductance of ∼20 pS (Figure 1D, Supplementary Figure 2B). These values matched those reported previously in cellular recordings^1^. At –100 mV, CNGA1 remained partially active (Pₒ ∼0.6), confirming the weak voltage dependence of this channel (Supplementary Figure 2B)^2^.

While CNGA1 also showed high activity with 200 µM cGMP, exhibiting Pₒ of 0.9–1.0, the incorporation of PIP2 markedly reduced activity: at 200 µM cGMP, 0.2% PIP2 lowered Pₒ by ∼40% (to ∼0.5), while 1.0% PIP2 suppressed channel opening by >90% (Pₒ <0.1) (Figure 1D). At saturating ligand (1 mM cGMP), inhibition was attenuated but still evident, with Pₒ reduced by ∼10% at 0.2% PIP2 and ∼20% at 1.0% PIP2. Importantly, the unitary conductance remained unchanged (∼20 pS in all conditions), indicating that PIP2 inhibition arises from altered gating by reducing single-channel open probability (Figure 1D). Together, these findings corroborate the ensemble flux assays and show that PIP2 reduces apparent cGMP sensitivity by reducing channel open probability.

To confirm that inhibition arises from the cytosolic leaflet, we analyzed CNGA1 single-channel activity with the soluble analog diC8-PIP2, which partitions into the inner leaflet and can be applied and washed off by perfusion. Application of 12.5 µM and 25 µM diC8-PIP2 decreased Pₒ in a dose-dependent manner, from ∼0.95 under control conditions to ∼0.6 and ∼0.1, respectively (Figure 1E). Furthermore, perfusion-mediated washout fully restored Pₒ to >0.9, demonstrating full reversibility of inhibition (Figure 1F). These results establish that PIP2 acts by binding directly to the cytosolic side of CNGA1 channels to suppress gating, consistent with the inhibitory effects of long-chain PIP2 observed in membranes and in agreement with our Tl⁺ flux and bilayer recordings.

### An intermediate gating state of CNGA1 revealed by cryo-EM in lipid nanodiscs

To gain mechanistic insight into PIP2 inhibition, we reconstituted purified CNGA1 channels into lipid nanodiscs (Supplementary Figure 1) and carried out single-particle cryo-EM analysis under multiple cGMP and PIP2 conditions. Nanodiscs were assembled using a lipid mixture of 65% DOPC, 11% POPE, and 24% POPS, which approximates the rod outer segment plasma membrane composition. To obtain PIP2-bound conformations, these lipids were supplemented with either 10% brain PIP2 or 3 mM diC8-PIP2. Datasets were collected in the absence (Supplementary Figures 3, 5, 7) and presence (Supplementary Figures 4, 6, 8) of saturating 3 mM cGMP.

In PIP2-free lipid environment, we obtained structures of CNGA1 in the expected closed conformation without cGMP, and the open conformation with cGMP (Supplementary Table 1). Both maps closely resembled the previously reported detergent-solubilized CNGA1 structures^6,16^. The nanodisc reconstructions, however, reached higher overall resolution (2.1–2.6 Å) than earlier detergent datasets (2.6–2.9 Å), permitting the visualization of new features. For example, in the nanodisc-embedded closed state, additional density consistent with a K⁺ ion was resolved immediately below the central gate, within the cytosolic vestibule (Supplementary Figure 9), suggesting that the cytosolic entryway may act as a K^+^ reservoir poised for permeation upon gate opening.

In addition, we also identified a previously unreported intermediate state under saturating cGMP conditions (Figure 2A, Supplementary Figure 4, and Supplementary Table 1). Roughly half of the particles (61.3%, ∼180k particles after all classifications and refinements) belonged to this class, with the remaining half forming the open state (Supplementary Figure 4). The intermediate conformation map was refined to 2.42 Å, enabling detailed interpretation. This state shared overall similarity with the closed conformation in its transmembrane domain and pore architecture, with the central gate sealed (Figure 2B–C). The CNBD of the intermediate conformation differs however from that of the closed state, showing changes that suggest partial activation. cGMP molecules were clearly resolved bound to the CNBD, and their occupancy correlated with a rotation and upward displacement (∼2.4 Å) of the CNBD C-helix relative to the closed state, although these changes are smaller than those occurring in the open state (Figure 2B). In the open state, the C-helix undergoes a larger displacement (∼6.6 Å), leading to a large upwards shift and rotation of the C-linker that stabilizes pore dilation. In the intermediate state, the C-linker exhibited only modest elevation relative to the closed state, which was insufficient to open the pore, leaving the conduction pathway occluded. Thus, the intermediate state is likely part of a gating transition in which cGMP binding has initiated conformational changes in the CNBD and C-linker, but were not fully transmitted to the pore.

**Figure 2.**
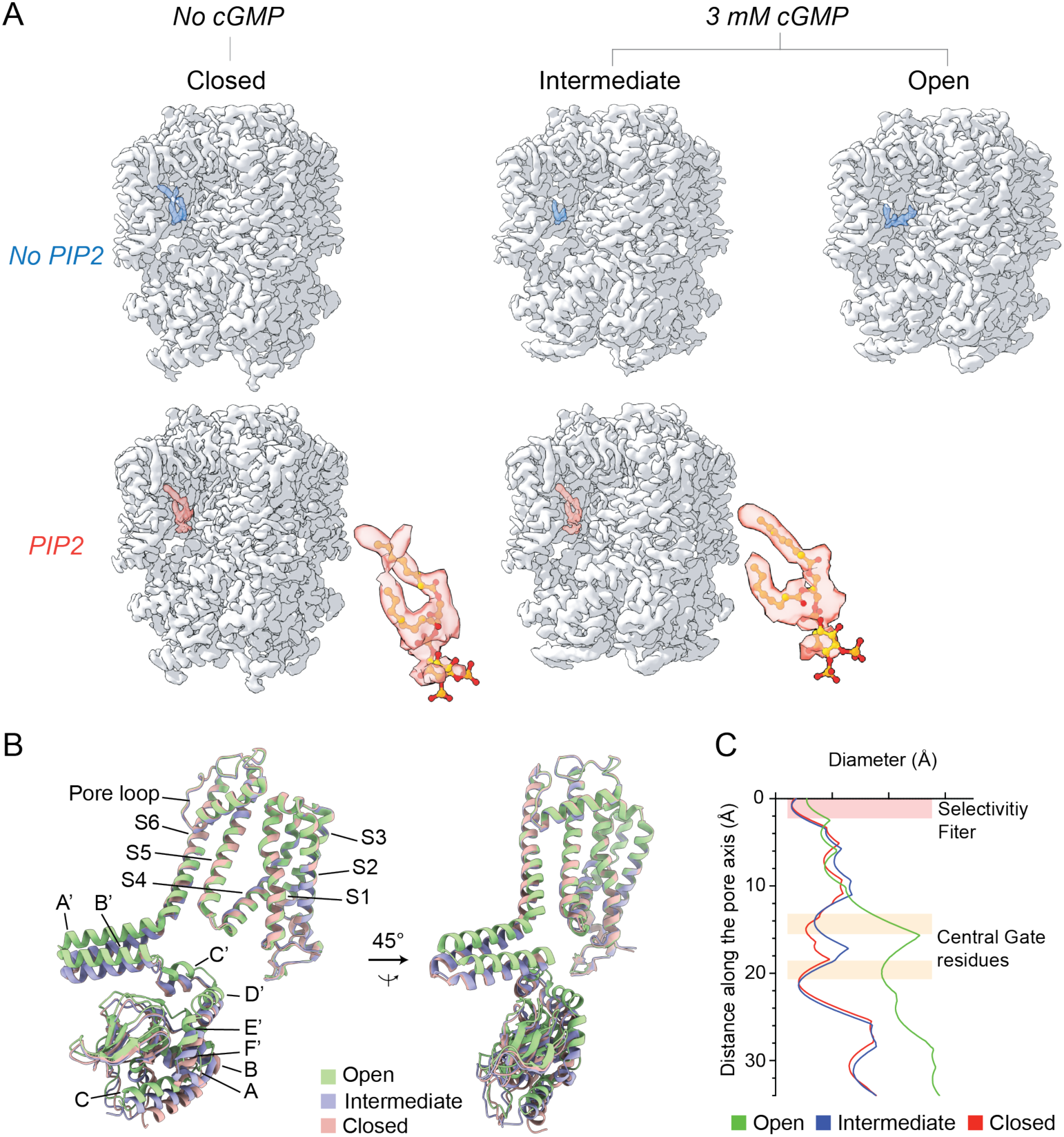
Cryo-EM structures of closed, intermediate, and open CNGA1 states with and without PIP2. (A) The top row shows three structures determined without exogenous PIP2, corresponding to the closed, intermediate, and open conformations. The bottom row presents the matching datasets collected in the presence of diC8-PIP2. Blue densities indicate lipid-like features consistently observed at the PIP2-binding pocket in PIP2-free structures, whereas red densities correspond to PIP2 in PIP2-supplemented reconstructions. PIP2 models (yellow) are overlayed in red densities and displayed at larger magnification. NB: no open conformation was observed with PIP2 and saturating cGMP. (B) Overlay of models for one CNGA1 subunit in closed (red), intermediate (blue), and open (green) conformations. (C) Pore diameters of the three states (closed, red; intermediate, blue; open, green). The selectivity filter and the central gate residues were indicated with red and beige-colored boxes, respectively.

### PIP2 binding prevents CNGA1 channel opening

We next examined CNGA1 channels reconstituted into nanodiscs containing 10% brain PIP2 (55% DOPC, 11% POPE, 24% POPS, 10% brain PIP2), and determined structures under apo and ligand-bound conditions (Supplementary Figures 5–6 and Supplementary Table 2). Strikingly, even in the presence of saturating cGMP (3 mM), no open channel conformation was detected (Supplementary Figures 6 and 10). Instead, particles exclusively populated the closed state in the absence of cGMP and the intermediate state in the presence of cGMP. Given that the intermediate represents a non-conducting form of the channel, these findings indicate that PIP2 shifts the conformational equilibrium away from the open state and stabilizes non-conductive conformations. This structural effect is fully consistent with our functional flux assays and single-channel recordings, which revealed robust inhibition of channel opening by PIP2. Notably, the overall architectures of CNGA1 in the closed and intermediate states were nearly indistinguishable from the corresponding conformations in PIP2-free nanodiscs, underscoring that PIP2 does not alter the structural framework of each state but instead biases their occupancy.

To determine where PIP2 binds we compared non-protein densities between intermediate-state structures obtained in the absence and presence of PIP2. In both datasets, non-protein densities were observed in inter-protomer grooves formed by the voltage-sensing domain (VSD), pore domain (PD), and C-linker. However, in the brain PIP2 dataset, imposition of C4 symmetry did not yield any detectable density corresponding to PIP2 headgroups in the inner leaflet region, likely because asymmetric/partial occupancies were averaged out. To recover such asymmetric densities, we applied symmetry expansion followed by two rounds of focused 3D classification using a mask covering one lipid-binding groove (Supplementary Figures 5–6). In both the closed and intermediate states, two of the four grooves consistently displayed a distinct enlarged head-group–like density, which could be fit by a PIP2 molecule, whereas the remaining two grooves contained only smaller densities consistent with generic fatty acyl chains. In the corresponding PIP2-free datasets, all four grooves displayed only acyl-chain–like densities without a detectable headgroup, indicating that PIP2 selectively occupies a subset of equivalent sites rather than saturating all four pockets (Figure 2A, top row, and Supplementary Figure 10).

Because the initial reconstructions using brain PIP2–reconstituted nanodiscs provided limited head-group detail, we also incubated CNGA1 nanodiscs prepared in the control lipid composition with a high concentration of soluble diC8-PIP2 (3 mM) to enhance visualization. Under this condition, we obtained the same closed and intermediate states only in the presence of PIP2 (with and without saturating 3 mM cGMP), and the reconstructions calculated with C4 symmetry revealed robust head-group–like densities bound at the inter-protomer grooves in both these states (Figures 2–3, Supplementary Figure 10, and Supplementary Table 2). These results validate phosphoinositide binding at the inter-protomer grooves and demonstrate that, when present in excess, PIP2 can occupy multiple equivalent sites on CNGA1.

**Figure 3.**
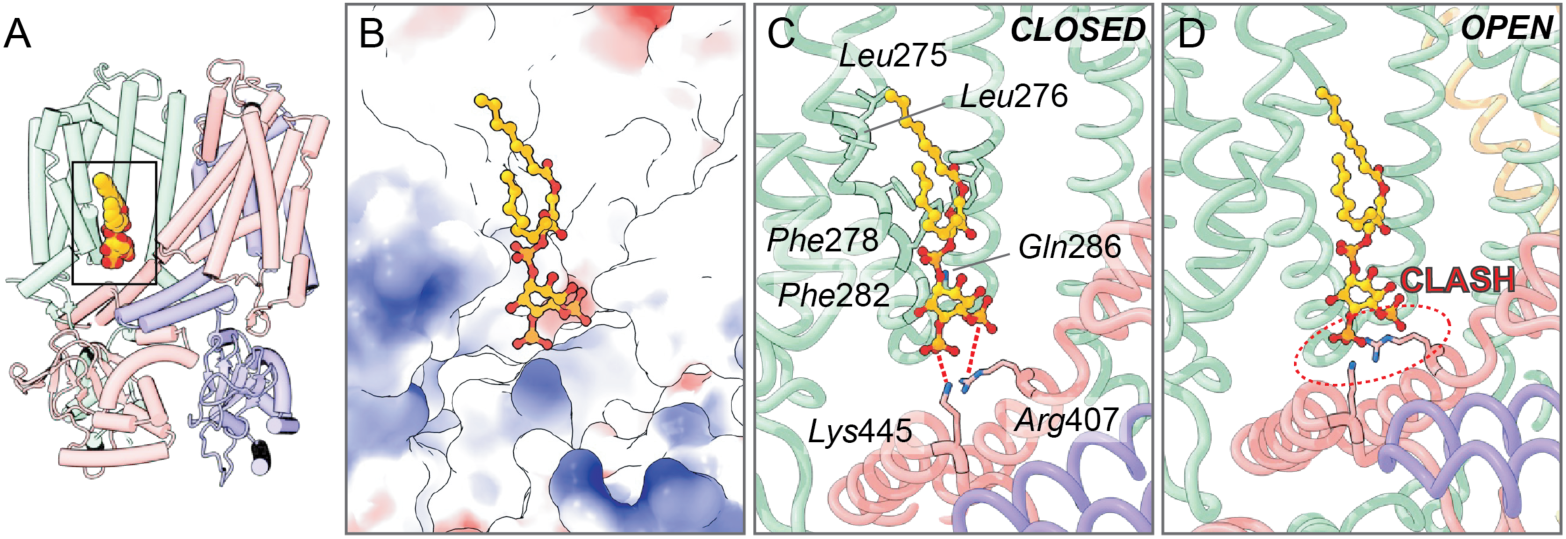
PIP2 binds at an allosteric site and prevents CNG channel opening. (A) Overview of the PIP2-bound closed CNGA1 conformation, highlighting the approximate PIP2 (yellow) binding site within the pocket surrounded by two adjacent subunits (red and green). (B) Electrostatic surface representation of the binding pocket showing negatively-charged PIP2 headgroup (red and yellow) residing in a positively charged cavity (blue). (C) Detailed binding mode of PIP2 in the closed state. The headgroup interacts electrostatically with positively charged side chains (red dashed lines) of Lys445 and Arg407 and is further stabilized by hydrophobic contacts along the acyl chains. (D) Superposition of the PIP2 molecule from the closed structure onto the open CNGA1 conformation, highlighting that the elevated C-linker in the open state sterically conflicts with the PIP2 headgroup (dashed red circle), resulting in a pronounced clash.

Notably, the positions of the PIP2-like densities were identical between the brain PIP2 and the diC8-PIP2 datasets across both the intermediate and closed conformational states, indicating that both brain and soluble PIP2 species occupy the same binding grooves. Accordingly, all subsequent structural analyses are based on CNGA1 structures determined in the presence of diC8-PIP2, enabling direct comparison with PIP2-free reconstructions at the same sites. In the closed state without PIP2, a well-defined density attributable to a fatty acyl chain was present within the groove, but no head-group density was detected, suggesting that the groove is not a PIP2-exclusive site but rather a permissive inner-leaflet lipid docking pocket that can be occupied by other phospholipids when PIP2 is absent. In contrast, in the open conformation, the non-protein densities are located elsewhere, in a deeper position within the groove near S6 and extended outward toward the region where the PIP2 headgroup resides in the PIP2-bound structures (Figure 2 and Supplementary Figure 10).

### PIP2 binding sterically obstructs C-linker elevation to prevent channel opening

Modeling the PIP2 density revealed that the two fatty acyl chains of PIP2 extended along a hydrophobic surface formed by the S4 and S5 helices (Leu275, Leu276, Phe278 and Phe282), while the inositol head group projected toward the cytosolic interface and localized near a positively-charged pocket formed by Gln286, Arg407, and Lys445 (Figure 3A-C). The positioning of the head group suggested electrostatic interactions with the guanidinium group of Arg407 and the amino group of Lys445 (Figure 3C). This orientation places the lipid head group directly above the C-linker, in the trajectory of the conformational elevation required for opening (Figure 3D).

To test the functional relevance of these putative contacts, we introduced alanine substitutions at Gln286, Arg407, and Lys445 and quantified channel activity in the presence of diC8-PIP2 (Figure 4A–C). Wild-type channels exhibited a robust decrease in Pₒ by >90% at 200 µM cGMP upon application of 25 µM diC8-PIP2. The Q286A mutant was inhibited by PIP2 similarly to wild type. The R407A mutant showed only ∼50% reduction under identical conditions, while the K445A substitution abolished inhibition, maintaining Pₒ >0.85 in the presence of diC8-PIP2 (Figure 4). Across all constructs, unitary conductance remained ∼20 pS, confirming that PIP2 alters gating transitions rather than unitary conductance (Figure 1, 4 and Supplementary Figure 2). These data establish that Arg407 and Lys445 are critical determinants of PIP2-mediated inhibition, with Lys445 playing the dominant role.

**Figure 4.**
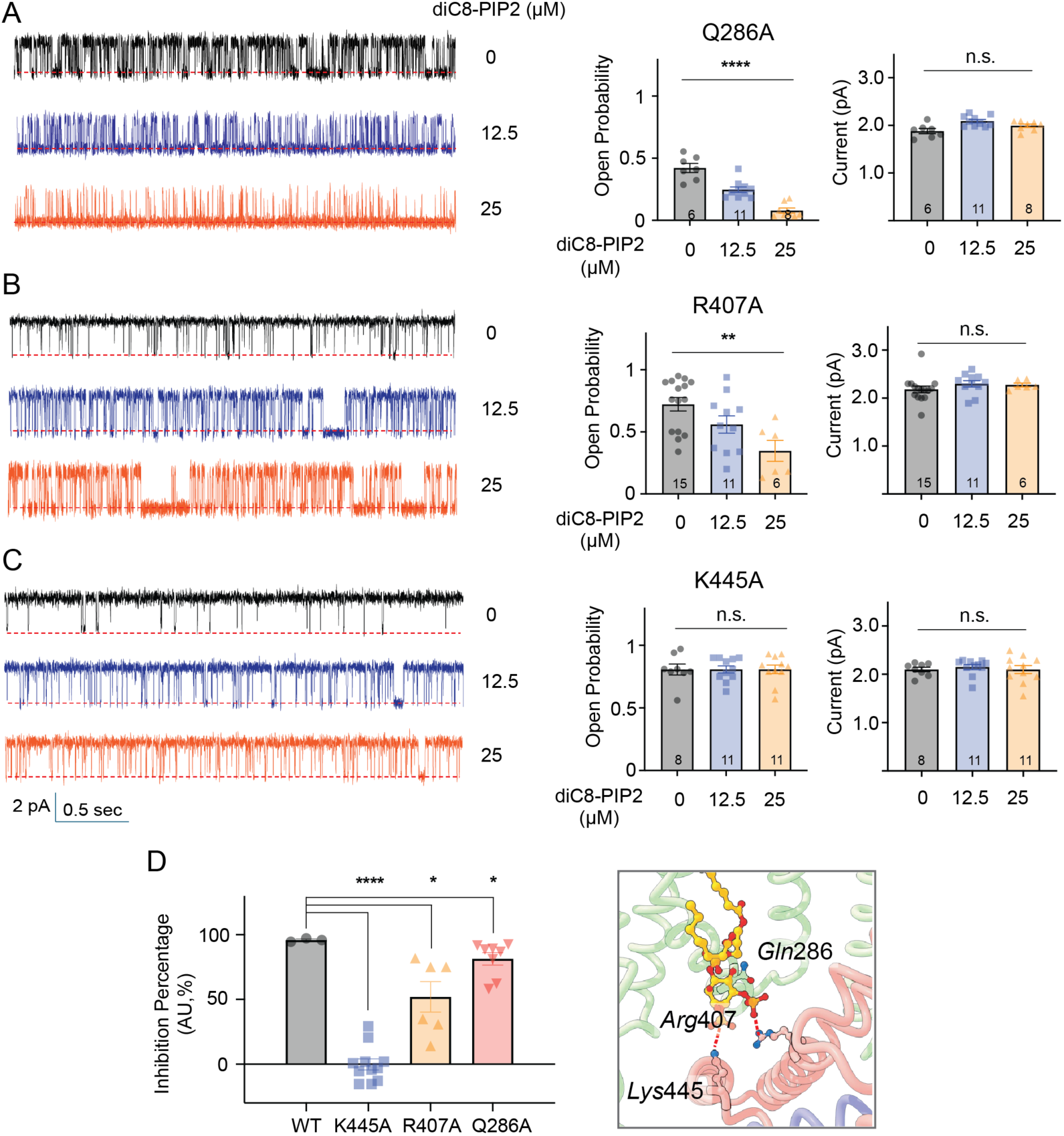
Validation of the CNGA1 PIP2-binding site using mutagenesis and single-channel electrophysiology. (A) Q286A; (B) R407A; (C) K445A. Left, representative single-channel traces, middle, open probability, right, unitary current amplitude at indicated diC8-PIP2 concentrations. (D) Percentage Inhibition of CNGA1 channel by PIP2 for WT and mutants with 25 μM diC8-PIP2. On the right, a detailed view of the PIP2 binding site in the closed CNGA1 indicating the mutated residues. Each data point was normalized to its maximal response and converted to an inhibitory scale as ‘100-100*(value/value_Max_)’. Traces were at +100 mV. The baseline is indicated by a red dotted line. Bars show mean ± SEM; each dot represents one independent bilayer recording. Statistics: within each mutant, conditions were compared by one-way ANOVA followed by Tukey’s multiple-comparison test in (A–C); ** p < 0.01, ****p < 0.0001, n.s. = not significant. The p values of Po in (A) and (B) were 5.8 × 10^-8^, 4.4 × 10^-3^, respectively. The inhibition percentage by diC8-PIP2 was compared by unpaired t-test (Welch’s) in (D); *p < 0.1, ****p < 0.0001. The p values for K445, R407, and Q286A mutants were 3.1 × 10^-10^, 1.3 × 10^-2^, and 1.9 × 10^-2^, respectively.

We next examined how interactions with these residues might interfere with gating motions. Superposition of the PIP2 model obtained from the PIP2-bound closed and intermediate structures onto the open-state model obtained in the absence of PIP2 revealed steric clashes between the elevated C-linker helix and the head group of PIP2 (Figure 3D). In the open state, upward displacement of the C-linker is necessary to propagate CNBD rearrangements into pore dilation, but in the presence of PIP2 this movement is sterically obstructed by bound PIP2. Arg407, positioned at the very end of the S6 helix, and Lys445, located in the C-linker, thus form a PIP2-binding site that anchors the gating ring in place and prevents its elevation. This steric constraint rationalizes the observed shift in cGMP sensitivity: cGMP binding to CNBD likely remains intact, but the conformational coupling between the CNBD and the gate is hindered, locking the channel in closed or intermediate states (Figure 3C–D). Collectively, these results demonstrate that PIP2 inhibits CNGA1 by directly engaging Arg407 and Lys445 to impede C-linker elevation, thereby stabilizing non-conductive conformations and preventing the transition to the open pore. This mechanism unifies the functional observation of reduced apparent cGMP sensitivity with the structural evidence of PIP2 densities in inter-protomer grooves, providing a detailed molecular model of phosphoinositide inhibition.

## Discussion

In this study, we defined the molecular mechanism by which PIP2 inhibits human rod CNGA1 channels. Ion flux assays and single-channel recordings established that PIP2 reduces the apparent cGMP sensitivity of purified CNGA1 without altering its unitary conductance in agreement with previous functional studies in cells^23^. To uncover the structural basis of this inhibition, we determined cryo-EM structures of nanodisc-reconstituted CNGA1 channels in the presence and absence of PIP2. This analysis revealed a previously uncharacterized intermediate conformation, providing additional structural detail for the cGMP-driven gating transition. Moreover, the structures showed that PIP2 prevents pore opening, even under saturating cGMP conditions, by locking the channel in closed and intermediate conformations. Focused classification and modeling identified PIP2 binding within inter-protomer grooves, where its head group forms interactions with key residues, R407 and K445. Mutational analysis confirmed the functional importance of these interactions, with K445 playing a dominant role. Structural superposition indicated that PIP2 binding inhibits channel opening by sterically hindering the upward movement of the C-linker, a conformational change needed to couple cGMP binding to the CNBD to pore opening. By restricting this motion, PIP2 effectively raises the energetic threshold for activation, requiring higher concentrations of cGMP to achieve channel opening. Thus, the reduction in apparent cGMP sensitivity can be directly attributed to PIP2 binding, which stabilizes closed and intermediate states by obstructing the CNBD–C-linker–to-gate coupling that drives pore gating (Figure 5A).

**Figure 5.**
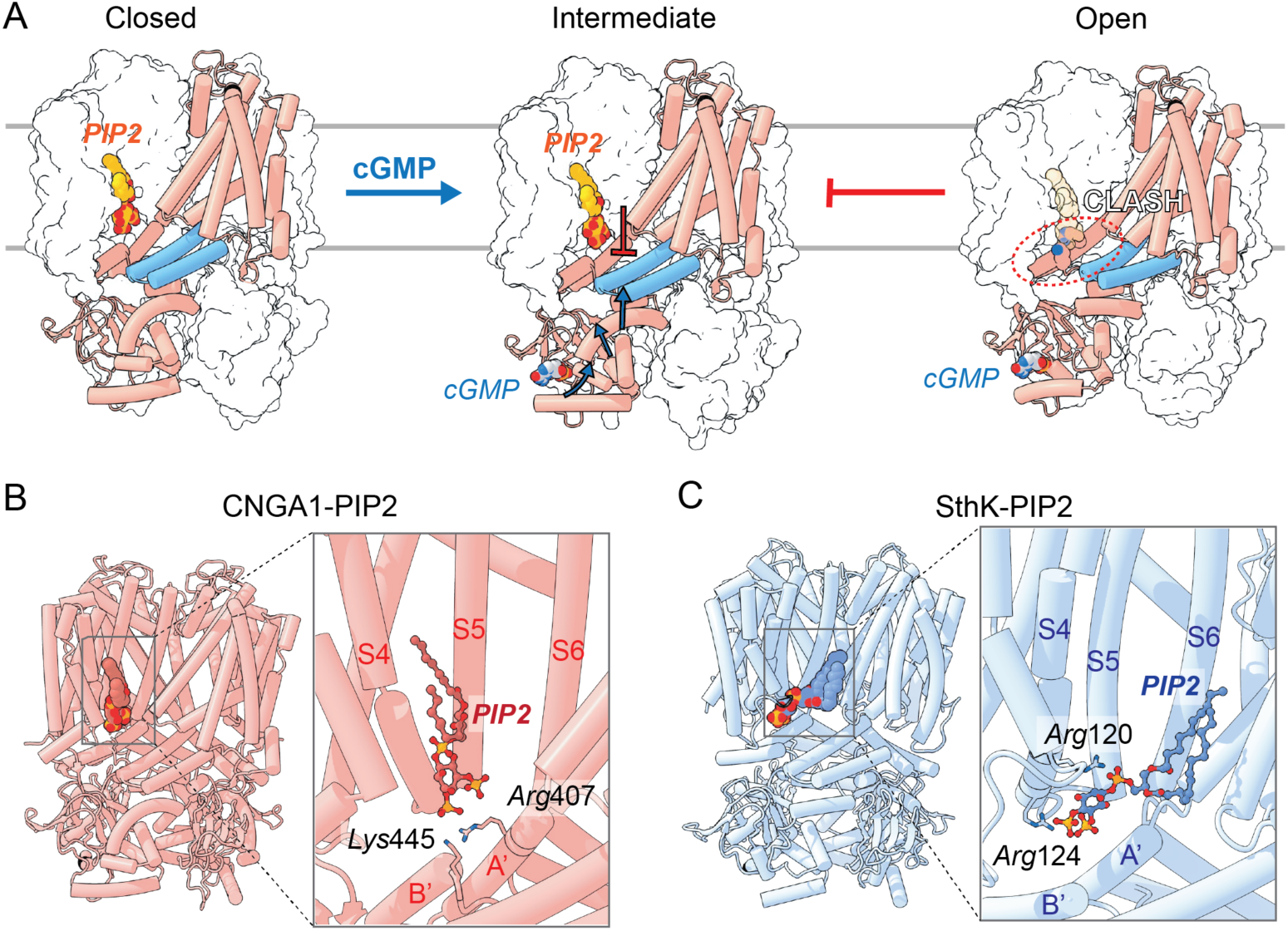
The mechanism of PIP2 inhibition in rod CNGA1 channels. (A) Cartoon of PIP2 inhibition mechanism of CNGA1 channels. The model is shown as red tubes for one subunit, with blue tubes indicating the adjacent subunit’s C-linker. PIP₂ is colored red-orange and cGMP blue-gray. The remaining regions are displayed as a white surface representation. (B–C) Comparison of PIP2 binding modes in CNGA1 (red) and SthK (blue, PDB ID:8VT9) channels with PIP2 binding highlighted.

Since an intermediate conformation of rod CNGA1 was not detected in previous detergent-solubilized preparations^16^ but was observed here in nanodisc-reconstituted samples (Figure 2A), one may hypothesize that the lipidic environment of nanodiscs might stabilize such a gating intermediate. However, intermediate-like ligand-bound conformations have also been reported for cone CNGA3/CNGB3 even under detergent conditions, indicating that the existence of such intermediates is not inherently dependent on nanodisc stabilization^35^. Rather, our results indicate that, for rod CNGA1, nanodisc reconstitution provides experimental conditions under which such intermediates become detectable, without implying that lipids are required to stabilize them. These observations highlight the value of lipid nanodiscs in resolving transitional conformations along the activation pathway of CNG channels and provide a structural framework for understanding how modulators such as PIP2 may bias the gating equilibrium.

Because these conclusions were derived from a homomeric CNGA1 background, an important question is whether the same inhibitory mechanism applies to the native rod channel. Although our structures were determined using a homomeric CNGA1 channel, native rod CNG channels assemble as a 3:1 CNGA1/CNGB1 heterotetramer^6^. The residues critical for PIP2 inhibition in CNGA1 (R407 and K445) are not conserved in CNGB1 (Supplementary Figure 11A). However, three of the four inter-subunit pockets still contain a CNGA1 protomer, and both the overall pocket geometry and the positive surface features that support PIP2 binding are well preserved in the heterotetramer (Supplementary Figure 11B). Furthermore, because CNG gating arises from inter-subunit–coupled conformational changes, modulation at these three conserved CNGA1-containing pockets would be expected to strongly influence the overall gating behavior of the heterotetramer, consistent with a similar PIP2-dependent inhibitory effect.

This work represents the first structural elucidation of PIP2 modulation in human CNG channels. Because phosphoinositides have also been reported to modulate the gating of other CNG/HCN family members—for example, PIP2 and PIP3 inhibition of cone CNG channels, PIP3 inhibition of olfactory channels, or PIP2 facilitation of HCN channel opening^27,30^—it is likely that similar mechanistic principles extend across this channel family. Comparisons with the bacterial CNG-like channel SthK support this view: as in CNGA1, where PIP2 engages C-linker residues R407 and K445 to impede the elevation of the C-linker required for opening, in SthK, PIP2 interacts with S4 residues R120 and R124 to also sterically inhibit C-linker elevation^31^. Although the detailed binding modes are not exactly the same, the site of action and the underlying molecular mechanism are broadly conserved (Figure 5B–C). Consistent with this idea, sequence alignment shows that the key basic residues corresponding to R407 and K445 are conserved in the principal cone (CNGA3) and olfactory (CNGA2) subunits, suggesting that these channels likely share the same phosphoinositide-dependent inhibitory mechanism (Supplementary Figure 11A). In contrast, HCN1–4 do not conserve these key residues, implying that phosphoinositides may act through alternative elements, potentially via VSD-linked interactions as observed in SthK, although this remains to be tested. Together, these observations provide a molecular framework for how phosphoinositides tune gating across the CNG and HCN superfamily and stabilize non-conducting states.

In addition to defining a mechanism for PIP2-dependent inhibition, our structures identify an allosteric inhibitory site within an inter-protomer lipid pocket that directly interfaces with the CNBD–C-linker–gate machinery. By occupying this pre-existing lipid-exposed groove, PIP2 shifts the conformational equilibrium toward non-conducting states, establishing this cavity as a bona fide allosteric inhibitory site in rod CNG channels. More broadly, lipid-facing pockets in membrane proteins, including ion channels and GPCRs^36–39^, are increasingly recognized as shared docking sites for endogenous lipids and synthetic small molecules and therefore represent privileged hot spots for allosteric regulation. In this context, the PIP2 pocket defined here provides a structurally resolved framework for designing modulators that mimic or antagonize phosphoinositide binding, offering a route to selectively tune CNG/HCN channel activity for future therapeutic targeting.

Beyond its structural and mechanistic implications, these findings carry important physiological significance. In rod photoreceptors, CNG channel activity determines the dark current and thereby sets the resting membrane potential, Ca²⁺ influx, and neurotransmitter release. By lowering the apparent cGMP sensitivity, PIP2 imposes an additional layer of control over channel gating, ensuring that only sufficiently high cGMP concentrations can trigger pore opening. Such inhibition would suppress background channel activity, limit noise in the dark state, and sharpen the fidelity of light responses. Moreover, PIP2 has already been implicated in multiple aspects of rod photoreceptor signaling, including light and dark adaptation and the regulation of phototransduction proteins such as transducin and PDE6^24–26,40,41^. Our findings extend this repertoire by identifying a new mode of modulation, in which PIP2 directly binds to and inhibits rod CNG channels, thereby reducing apparent cGMP sensitivity. Together, this work highlights a mechanistic paradigm by which phosphoinositides bias ligand-gated channels toward non-conducting conformations and reshape their gating behavior, underscoring their central role as versatile lipid regulators across the CNG/HCN channel superfamily.

## Materials and Methods

### Protein expression and purification

The truncated human CNGA1 construct (residues 145–690, N-terminal Flag tag in pEZT-BM) was kindly provided by Dr. Youxing Jiang (UT Southwestern Medical Center). Protein expression and purification were performed following the method reported by the Jiang lab^16^, with several modifications as detailed below.

Bacmids were generated in *E. coli* DH10Bac and baculoviruses were amplified in Sf9 cells using Cellfectin II (Thermo Fisher Scientific). For protein production, HEK293F cells (Thermo Fisher Scientific) were infected at a virus-to-cell ratio of 1:10 (v/v) and supplemented with 10 mM sodium butyrate 16 h after the baculovirus infection, followed by 48 h incubation at 37 °C. Harvested cells were resuspended in purification buffer (25 mM HEPES, 150 mM KCl, pH 7.4) containing 50 μM cGMP and a tablet of protease inhibitors cocktail, cOmplete (Roche), and membranes were solubilized in 1% (w/v) lauryl maltose neopentyl glycol (LMNG) (Anatrace) for 1.5 h. After centrifugation, the supernatant was incubated with anti-DYKDDDDK G1 affinity resin (Genscript) for 1 h, washed with the purification buffer containing 0.06% glyco-diosgenin (GDN) (Anatrace) and 50 μM cGMP, and eluted with 0.15 mg/ml Flag peptide (MedChemExpress). The eluate was concentrated and subjected to size-exclusion chromatography on a Superose 6 Increase 10/300 GL column (Cytiva) in the purification buffer with 0.06% GDN and 50 μM cGMP. Peak fractions were pooled and concentrated to 10 mg/ml with 1 Abs_280_ = 1 mg, and subsequently reconstituted into proteoliposomes for functional assays and into nanodiscs for cryo-EM analysis. All purification steps were performed at 4 °C (Supplementary Figure 1A).

Site-directed mutagenesis was used to generate the Q286A, R407A, and K445A substitutions (Q5 polymerase, NEB). All CNGA1 mutants were produced and purified following the established protocol, yielding comparable levels of protein with similar purity.

### Reconstitution of channels into liposomes for functional assays

For Tl⁺ stopped-flow experiments, purified CNGA1 channels were incorporated into LUVs preloaded with the fluorophore ANTS (8-aminonaphthalene-1,3,6-trisulfonic acid; Life Technologies), following the protocol reported previously^19,32^. Lipid mixtures (5 mg total; 65:11:24 molar ratio of DOPC:POPE:POPS, with or without added PIP2) were dried to a thin film under nitrogen gas and subsequently desiccated overnight under vacuum. The resulting lipid film was hydrated with pre-mix buffer (10 mM HEPES, 140 mM KNO₃, pH 7.4) and solubilized by sonication in 33 mM CHAPS. Protein was incorporated at a ratio of 30 µg per mg lipid, and ANTS dye was added during reconstitution. An initial aliquot of ANTS stock (75 mM in the pre-mix buffer, pH 7.4) was combined with the lipid–protein mixture and incubated at room temperature for 20 min, after which additional ANTS was introduced to reach a final concentration of 25 mM. Detergent was removed by sequential incubations with SM-2 Bio-Beads (Bio-Rad; 330 mg from a 50% slurry in the pre-mix buffer), first for 2 h and then 15 h with a fresh batch at 21 °C under gentle agitation. The supernatant containing liposomes was collected, briefly sonicated in a water bath, and passed through a 0.1 µm polycarbonate membrane using a mini-extruder (Avanti Polar Lipids). Excess extravesicular ANTS was removed via a PD-10 desalting column (GE Healthcare). To assess reconstitution efficacy, proteoliposomes containing 0, 0.2, 1.0, and 2.5% brain PIP2 were analyzed by SDS-PAGE, confirming comparable protein incorporation across conditions (Supplementary Figure 1B). Prior to stopped-flow measurements, vesicle suspensions were diluted in the pre-mix buffer to a final volume of 22 mL for single-mixing recordings.

For planar bilayer single-channel recordings^42–44^, liposomes were generated from a lipid mixture (5 mg total; 65:11:24 molar ratio of DOPC:POPE:POPS, with or without PIP2). Lipids dissolved in chloroform were dried under a continuous stream of nitrogen in glass tubes to produce a thin film and subsequently rehydrated with reconstitution buffer (10 mM HEPES, 400 mM KCl, 5 mM NMDG, pH 7.6). The suspension was solubilized with 33 mM CHAPS by brief sonication in a water bath. Purified CNGA1 protein was incorporated at 40 µg per mg of lipid, and the mixture was incubated for 20 min at room temperature. Detergent removal and liposome formation were achieved by gravity flow through an 18 ml Sephadex G-50 column (GE Healthcare), with eluate collected in 500 µl fractions. Fractions containing liposomes were identified by turbidity, pooled, and divided into 70 µl aliquots. Samples were flash frozen in liquid nitrogen and stored at –80 °C until use.

### Tl⁺ stopped-flow assay

Stopped-flow fluorescence measurements were performed at 25 °C using a SX.20 LED spectrophotometer (Applied Photophysics, ProDataSX software, Leatherhead, UK) in single-mix mode. Excitation was set to 360 nm, and emission was monitored with a 420 nm long-pass filter. Prior to mixing with quencher, LUVs containing reconstituted protein were incubated with increasing concentrations of cGMP (8–2500 µM). Samples were then rapidly mixed 1:1 with quenching buffer (10 mM HEPES, 90 mM KNO₃, 50 mM TlNO₃, 200 µM cGMP, pH 7.4), and ANTS fluorescence decay was recorded for 1 s at a sampling density of 5000 points. Each condition was measured in eight technical replicates. Control traces were collected using reconstitution buffer lacking quencher or cGMP. All experiments were independently repeated at least three times with separately prepared proteoliposomes.

Data analysis was performed in Matlab following established procedures^19,32,45^. Because vesicle size heterogeneity and variable channel content per liposome broadened the fluorescence decay, the first 100 ms of each trace were fitted to a stretched exponential (Eq. 1). The fitted parameters were subsequently used to calculate Tl⁺ influx rates at 2 ms (Eq. 2).

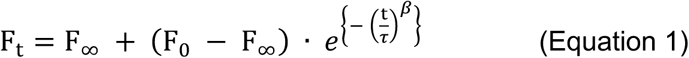

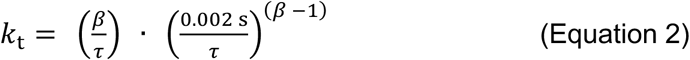

Here, Fₜ, F₀, and F∞ denote fluorescence at time t, the initial fluorescence, and the asymptotic final value, respectively; τ is the characteristic time constant, β is the stretching exponent, and kₜ represents the calculated influx rate (s⁻¹) at 2 ms. For each experimental repeat, paired measurements were performed with and without PIP2 using aliquots from the same stock solutions. Flux rates obtained in the presence of PIP2 were normalized to the corresponding no-PIP2 control to account for preparation variability.

### Planar bilayer single-channel recording

Planar lipid bilayers were formed in a horizontal setup, where the two chambers were separated by a partition containing a 100 µm aperture. Bilayers were generated by spreading a small air bubble containing DPhPC (1,2-diphytanoyl-sn-glycero-3-phosphocholine, Avanti Polar Lipids; 8 mg/ml in decane) across the aperture using the tip of a 10 µl pipette. Successful bilayer formation was verified by capacitance monitoring, and only bilayers within the range of 25–50 pF were used for recordings. Both chambers were filled with filtered recording solution (100 mM KCl, 10 mM HEPES, pH 7.4). To ensure unidirectional channel activation, 40, 200, and 1000 µM cGMP was added exclusively to the bottom chamber, thereby permitting activity only from channels oriented with their cyclic nucleotide–binding domains facing this side (Supplementary Figure 2A).

For phosphoinositide perfusion experiments, diC8-PIP2 was introduced into the bottom chamber (the cGMP-containing side), and currents were recorded at both +100 mV and −100 mV in the presence of 200 µM cGMP. Following PIP2 exposure, the bottom chamber was washed by perfusing ∼2 mL of PIP2-free recording solution to remove residual diC8-PIP2.

Currents were recorded with an Axopatch 200B amplifier (Molecular Devices) and digitized at 20 kHz using a Digidata 1440A interface, with pClampex 10.7.0.3 software (Molecular Devices) controlling acquisition. An eight-pole Bessel filter was applied online at 2 kHz. Single-channel events were analyzed with pClampfit 10.7.0.3 (Molecular Devices). Amplitude histograms were normalized to the total number of events such that the cumulative peak heights summed to 1.

For statistical analyses, a one-way ANOVA followed by Tukey’s multiple-comparison test was employed with a 95% confidence level. Significance thresholds were assigned as follows: P < 0.05 (*), P < 0.01 (**), P < 0.001 (***), and P < 0.0001 (****).

### Reconstitution of channels into nanodiscs and data collection for cryo-EM

For cryo-EM analysis, CNGA1 channels were incorporated into lipid nanodiscs^46^ (Supplementary Figure 1C). Three different lipid environments were examined: (1) a control mixture of 65% DOPC, 11% POPE, and 24% POPS, (2) the same mixture supplemented with 10% PIP2 (replacing DOPC), and (3) the control mixture with 3 mM soluble PIP2 added. Lipid films were prepared by drying the chloroform-dissolved lipid mixtures under a stream of nitrogen, followed by overnight desiccation under vacuum. The dried films were resuspended in reconstitution buffer (25 mM HEPES, 150 mM KCl, pH 7.4) and solubilized with 200 mM sodium cholate. The resulting lipid stocks were flash frozen in liquid nitrogen and stored at –80 °C until use. Membrane scaffolding protein MSP1E3 was expressed and purified according to standard protocols, concentrated to 10 mg/ml in reconstitution buffer, and mixed with purified CNGA1 (10 mg/ml after gel filtration). CNGA1, MSP1E3, and lipids were combined at a molar ratio of 1:1.1:70 in reconstitution buffer, yielding a final cholate concentration of 15 mM. After incubation for 20 min at room temperature, detergent was removed by sequential incubations with pre-washed SM-2 Bio-Beads (Bio-Rad): first for 2 h and then overnight (15 h) with fresh beads under gentle agitation. The cleared supernatant was collected, and Bio-Beads were washed with 250 µl reconstitution buffer to yield a final sample volume of 500 µl. Following filtration through a Spin-X column, nanodiscs containing tetrameric CNGA1 were separated on a Superose 6 10/300 column (Cytiva) equilibrated with reconstitution buffer. Peak fractions were pooled and concentrated to 8 mg/ml (A280 conversion: 1 OD = 1 mg/ml). For cryo-EM grid preparation, the final sample was adjusted to 7 mg/ml CNGA1 in reconstitution buffer supplemented with 3 mM fluorinated Fos-choline-8 (Anatrace), either with or without 3 mM cGMP. In total, six CNGA1–nanodisc samples were generated: three lipid compositions prepared in the presence or absence of cGMP (Supplementary Figure 1C).

Cryo-EM grids were prepared under identical conditions. Gold UltrAuFoil R1.2/1.3 300 mesh grids (Quantifoil) were glow discharged (PELCO easiGlow, 80 s, –25 mA) and loaded with 3 µl of sample. After incubation for 20 s at 22 °C and 100% humidity, grids were blotted for 2 s (blot force 0) and plunge frozen in liquid ethane using a Vitrobot Mark IV (FEI, Thermo Fisher). Initial grid screening was performed on a 200 kV Glacios microscope (Thermo Fisher) to optimize ice thickness, particle distribution, and preferred orientation before high-resolution data collection on the 300 kV Titan Krios.

Data were collected on a Titan Krios G3i microscope operated at 300 kV, equipped with a Gatan K3 direct electron detector and a BioQuantum energy filter (20 eV slit width), controlled by Leginon^47^. Movies were acquired in super-resolution mode at 105,000× nominal magnification, corresponding to a pixel size of 0.4125 Å. Each movie consisted of 40 frames recorded at 45 ms per frame, with exposure time of 1800 ms. For CNGA1 reconstituted into nanodisc containing brain PIP2 (55% DOPC, 11% POPE, 24% POPS, 11% brain PIP2) (in absence and presence of cGMP), an exposure rate was 26.98 e-/Å2/s, corresponding to a total dose of 48.56e⁻/Å². Defocus range was 0.6–2.5 µm using Leginon. For CNGA1 reconstituted into nanodisc containing the control lipid composition (65% DOPC, 11% POPE, 24% POPS) (in absence and presence of cGMP), an exposure rate was 26.92 e-/Å2/s, corresponding to a total dose of 48.45e⁻/Å². Defocus range was 0.5–3.0 µm using Leginon. For CNGA1 reconstituted into nanodisc containing the control lipid composition (65% DOPC, 11% POPE, 24% POPS) incubated with diC8-PIP2 (in absence and presence of cGMP), an exposure rate was 26.93 e-/Å2/s, corresponding to a total dose of 48.48e⁻/Å². Defocus range was 0.6–2.4 µm using Leginon.

### Processing of Cryo-EM data

For all the samples, cGMP-free and cGMP-bound CNGA1 reconstituted into MSP1E3 nanodiscs, containing various lipid compositions, 1) Control (65% DOPC, 11% POPE, 24% POPS), 2) Brain PIP2 (55% DOPC, 11% POPE, 24% POPS, 10% brain PIP)_2_, 3) DiC8-PIP2 (65% DOPC, 11% POPE, 24% POPS in presence of 3 mM diC8-PIP2) datasets, the processing was done in Relion 5.0-beta3^48^.

For cGMP-free CNGA1 in the PIP2-free nanodiscs dataset (Supplementary Figure 5), 8718 movies were collected in super-resolution mode, dose weighted and binned by 3 using MotionCor2^49^ in Relion 5^48^. The contrast transfer function (CTF) on the resulting dose-weighted micrographs were determined by CTFFIND4^50^. 1.48 million particles were picked by CrYolo 1.8.0^51^ and extracted with a box size of 256 pixels. After 2 rounds of 2D classification, 0.82 million particles were selected for 3D initial model in Relion for the initial map reference. These particles were subjected to 3D classification with alignment, without symmetry into 6 classes, using the initial map lowpass-filtered to 60 Å as a reference. The classes without density or low-resolution density in transmembrane region were discarded. 4 classes with 0.71 million particles were selected and the orientation and translational parameters for these selected particles were refined with the auto-refine algorithm of Relion without imposing symmetry, using the initial map lowpass-filtered to 40 Å as a reference and a mask surrounding the protein and excluding the nanodisc density. The resulting refined particle images were subjected to 3D classification without image alignment. The classes without density or low-resolution density in transmembrane region were discarded. Three classes with 0.70 million particles were selected. The particles in these classes which had no difference between them, subjected to 3D refinement in 15 Å-low pass filter, CTF refined and polished with Bayesian polishing^52^ to get a map with a resolution of 2.56 Å. Because any difference between each protomer was not detected, the map was applied C4 symmetry with the initial map lowpass-filtered to 15 Å as reference to generate a map with resolution of 2.46 Å. And the orientation and translational parameters for these particles were unbinned with a box size of 384 pixels subjected to 3D refinement, CTF refined and polished with Bayesian polishing to get a map with a resolution of 2.07 Å. After unbinning, all subsequent 3D refinements were performed with Sidesplitter to prevent overfitting of the maps. This class is referred to as the control closed state because the there was no density for cGMP at CNBD, the C helix was not elevated, and the pore gate was tightly closed.

For cGMP-bound CNGA1 in the control nanodiscs dataset (Supplementary Figure. 6), 7827 movies were collected in super-resolution mode, dose weighted and binned by 3 using MotionCor2 in Relion. The contrast transfer function (CTF) on the resulting dose-weighted micrographs were determined by CTFFIND4. 1.49 million particles were picked by CrYolo 1.8.0 and extracted with a box size of 256 pixels. After two rounds of 2D classification, 0.58 million particles were selected for 3D initial model in Relion for the initial map reference. These particles were subjected to 3D classification with alignment, without symmetry into 6 classes, using the initial map lowpass-filtered to 60 Å as a reference. The classes without density or low-resolution density in transmembrane region were discarded. Three classes with 0.42 million particles were selected and the orientation and translational parameters for these selected particles were refined with the auto-refine algorithm of Relion without imposing symmetry, using the initial map lowpass-filtered to 40 Å as a reference and a mask surrounding the protein and excluding the nanodisc density. The resulting refined particle images were subjected to 3D classification without image alignment. The classes without density or low-resolution density in transmembrane region were discarded. Two classes with 331 thousand particles and a class with 81 thousand particles were selected as class 1 and class 2, respectively. The particles of class 1 and class 2 were subjected to multiple rounds of 3D refinement in 15 Å-low pass filter and C4 symmetry, CTF refinement, Bayesian polishing and 3D refinement to generate maps with resolution of 3.02 and 3.33 Å. The map of one of classes from the 3D classification of the class 1, class 1-3, seems similar to that of the class 2. The particles of the class 1-3 and class 2 were pooled and subjected to multiple rounds of 3D refinement, CTF refinement, Bayesian polishing (unbinned with a box size of 384 pixels, after unbinning, all subsequent 3D refinements were performed with Sidesplitter to prevent overfitting of the maps.) and 3D refinement to generate maps with resolution of 2.62 Å. This class is referred to as the control open state because there were densities for cGMP at CNBD, the cytosolic domain including C helix was elevated, and the pore gate was fully open, which is almost identical to the structure of open structure of detergent-solubilized CNGA1 published before^16^.

The maps of the class 1-1 and class 1-2 were similar in terms of conformation but that of the class 1-1 has weaker lipid-like densities in the lipid binding pockets between protomers, while the map of the class 1-2 has relatively strong lipid-like densities in the pockets. Therefore, the two classes’ particles are subjected to multiple rounds of 3D refinement, CTF refinement, Bayesian polishing (unbinned with a box size of 384 pixels) and 3D refinement separately to get maps with a resolution of 2.99 Å and 2.45 Å. The class 1-2 is referred to as the control intermediate state because there were densities for cGMP at CNBD, the cytosolic domain including C helix was slightly elevated, and the pore gate was tightly closed, which has both features of closed and open states.

The cGMP-free and cGMP-bound CNGA1 in the brain PIP2 nanodiscs were processed in a similar manner as the cGMP-free CNGA1 in the control nanodiscs dataset (Supplementary Figure 7 and 8). For the cGMP-free and cGMP-bound datasets, 6002 / 7932 movies were collected in super-resolution mode and motion-corrected (bin 3, dose-weighted) in RELION, followed by CTF estimation with CTFFIND4. Particles were picked by CrYOLO (∼1.63 M / ∼1.32 M for the ligand dataset) and subjected to 2D classification, yielding ∼1.10 M / ∼0.82 M particles for initial 3D model generation. Subsequent 3D classification with alignment (C1) followed by classification without alignment removed low-quality classes, resulting in 0.83 M / 0.51 M particles for high-resolution refinement. These particles were refined with a low-pass–filtered reference, CTF-refined, and polished to obtain a 2.51 Å / 2.49 Å map. For the cGMP-free dataset, C4 symmetry was then imposed before unbinning, yielding 2.46 Å and, after unbinning and further refinement, 1.97 Å. In contrast, for the cGMP-bound dataset, C4 symmetry was imposed at the time of unbinning, which directly yielded a 2.21 Å map that did not improve further upon subsequent refinement (unbinned with a box size of 384 pixels). All final refinements were performed with Sidesplitter^53^ to minimize overfitting. Across the repeated 3D classification steps during refinement, the cGMP-free dataset consistently yielded a single closed conformation indistinguishable from previously reported closed structures. By contrast, the cGMP-bound dataset converged to a single intermediate conformation in which the pore remained closed, but cGMP density was clearly present at the CNBD and the cytosolic domains were elevated relative to the closed state, although not to the level seen in fully open structures.

Symmetry expansion followed by two consecutive rounds of 3D classification without alignment (τ = 20) was performed on both datasets using the same residue-defined local mask encompassing five helices forming the pocket (S4–S5, 268–318; S6, 376–406 of the target protomer, and S5, 291–317; S6, 382–405 of the adjacent protomer). The mask was generated separately from the closed-state model for the cGMP-free dataset and from the intermediate-state model for the cGMP-bound dataset. both datasets, these classifications were specifically examined to determine whether any class exhibited lipid density consistent with a PIP2-like headgroup near the inner leaflet.

Among the expanded particles in the cGMP-free dataset (2.6 M after symmetry expansion), 29.6% (0.78 M) fell into a class that displayed such a density with a large headgroup positioned near the inner leaflet. These particles were isolated and subjected to 3D refinement under C1, followed by CTF refinement and Bayesian polishing, yielding a 2.45 Å closed-state structure. Among the expanded particles in the cGMP-bound dataset (1.3 M after symmetry expansion), 9.5% (0.12 M) fell into a class that displayed such a density with a large headgroup positioned near the inner leaflet. These particles were isolated and subjected to 3D refinement under C1, followed by CTF refinement and Bayesian polishing, yielding a 2.62 Å intermediate-state structure.

The cGMP-free and cGMP-bound CNGA1 in nanodiscs with diC8-PIP2 were processed in a similar manner as the other datasets (Supplementary Figure 9 and 10). For the cGMP-free and cGMP-bound datasets, 7396 / 6835 movies were collected in super-resolution mode and motion-corrected (bin 3, dose-weighted) in RELION, followed by CTF estimation with CTFFIND4. Particles were picked by CrYOLO (∼1.25 M / ∼1.21 M for the ligand dataset) and subjected to 2D classification, yielding ∼0.74 M / ∼0.70 M particles for initial 3D model generation. Subsequent 3D classification with alignment (C1) followed by classification without alignment removed low-quality classes, resulting in 0.56 M / 0.43 M particles for high-resolution refinement. These particles were refined with a low-pass–filtered reference, CTF-refined, and polished to obtain a 2.48 Å / 2.66 Å map. For both the cGMP-free and cGMP-bound datasets, particles were first re-extracted unbinned (box size 384 px) and subsequently refined, after which C4 symmetry was imposed. This yielded 2.11 Å / 2.20 Å maps for the two datasets, respectively. All final refinements were performed with Sidesplitter to minimize overfitting. Across the repeated 3D classification steps during refinement, the cGMP-free dataset consistently yielded a single closed conformation, whereas the cGMP-bound dataset yielded a single intermediate conformation.

### Model building and validation

For model building, previously published CNGA1 detergent-solubilized structures (apo closed and open structures, PDB ID: 7LFT and 7LFW^16^, respectively) were placed into the density using UCSF Chimera^54^ and manually adjusted in COOT^55^. The intermediate structures modeled based on the closed CNGA1 structure (PDB ID: 7LFT). Models were real-space refined in Phenix^56,57^ and further improved using ISOLDE^58^. cGMP and lipids were manually added to each model in COOT and outliers were fixed. Quality of the final models was validated with MolProbity^59^ against unfiltermed half maps and the unsharpened full map. All structural figures were prepared using ChimeraX^60^.

## Data availability

The maps and models have been deposited at the Electron Microscopy Data Bank (EMDB) and the Protein Data Bank (PDB). Accession codes are: CNGA1 channel closed state in nanodisc cGMP-free (EMD-74534/9ZPV); CNGA1 channel intermediate state in nanodisc cGMP-bound (EMD-74535/9ZPW); CNGA1 channel open state in nanodisc cGMP-bound (EMD-74536/9ZPX); CNGA1 channel closed state in nanodisc with brain PIP2 cGMP-free (EMD-74537/9ZPY); CNGA1 channel intermediate state in nanodisc with brain PIP2 cGMP-bound (EMD-74538/9ZPZ); CNGA1 channel closed state in nanodisc with diC8-PIP2 cGMP-free (EMD-74539/9ZQ0); CNGA1 channel intermediate state in nanodisc with diC8-PIP2 cGMP-bound (EMD-74540/9ZQ1). Raw electrophysiology and fluorescence traces are available from the corresponding author upon request. Sample raw data traces are provided in the corresponding figures. Electrophysiology and functional data are not deposited due to their complexity and large size and requiring the use of proprietary scientific software to open and analyze. All data supporting the findings of this paper are available from the corresponding author upon request.

## Acknowledgements

The cryo-EM data for this study were collected at the Weill Cornell Medicine cryo-EM facility and the NYU Langone cryo-EM facility (RRID: SCR_019202). We thank Edwin C. Fluck of the Weill Cornell Medicine cryo-EM facility and Bing Wang, Huihui Kuang, and Bill Rice of the NYU Langone cryo-EM facility for help with screening and collecting cryo-EM data. We also thank Youxing Jiang (Department of Biophysics, University of Texas Southwestern Medical Center) for generously providing us with pEZT-BM-CNGA1 plasmid used for making CNGA1-expressing baculovirus. Some figures were drawn with BioRender.com. The work was supported by National Institutes of Health (NIH) grant, GM124451 to C.M.N.

## Author contributions

T.P. and C.M.N. designed this research; T.P. performed all experiments and analyzed the data under the supervision of C.M.N; T.P. and C.M.N wrote the manuscript.

## Declaration of interests

The authors declare no competing interests.

## Supplementary Figures

**Supplementary Figure 1.**
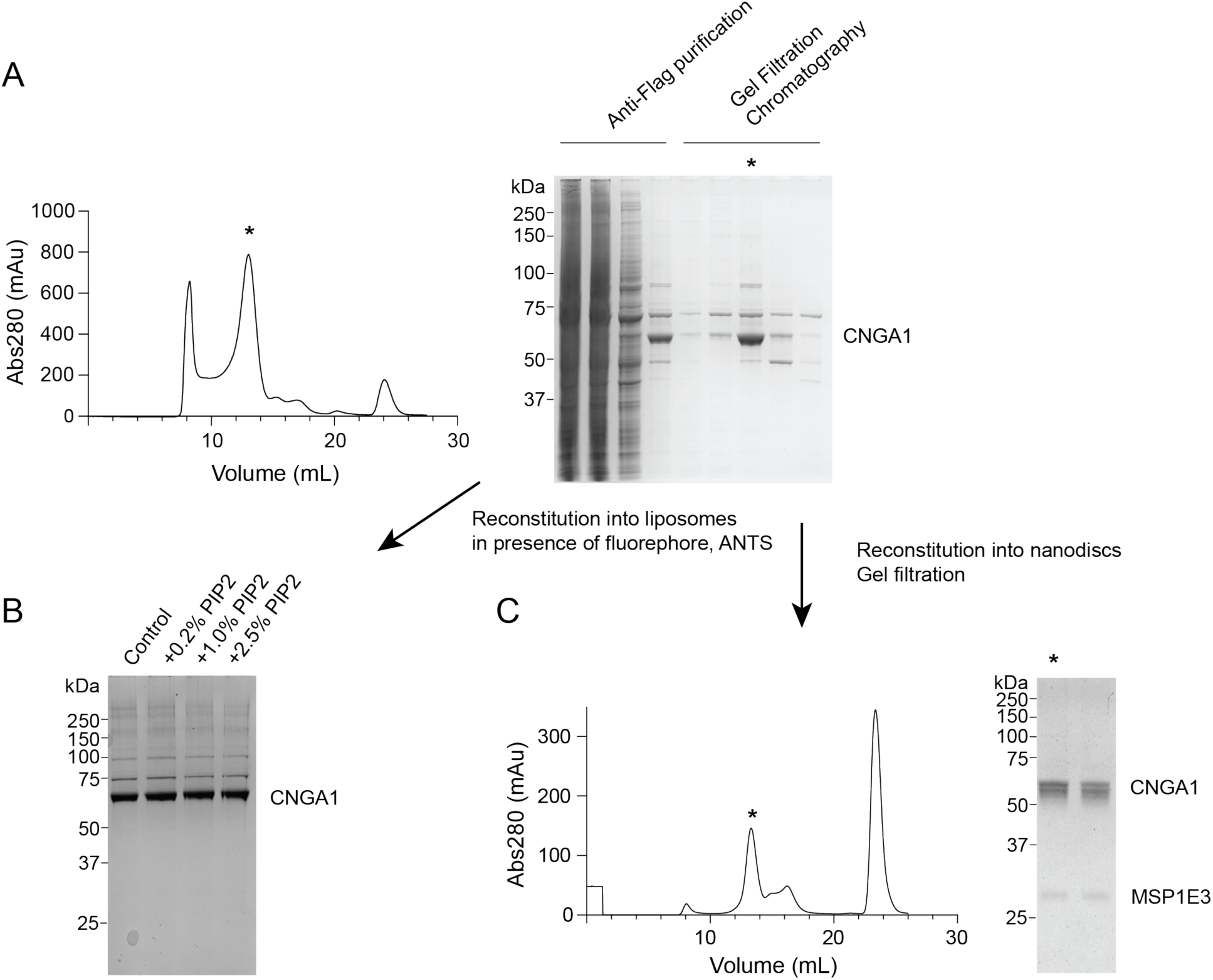
Purification and reconstitution of CNGA1. (A) Purification profile of CNGA1 by anti-Flag affinity purification and size exclusion chromatography (SEC). The peak corresponding to CNGA1 is indicated with asterisk (*). (B) SDS–PAGE analysis of CNGA1 after reconstitution into proteoliposomes containing different concentrations of PIP2 for Tl^+^ influx stopped-flow assay. (C) SEC profile of CNGA1 after reconstitution into nanodiscs. The major peak corresponding to reconstituted CNGA1 is indicated with asterisk (*).

**Supplementary Figure 2.**
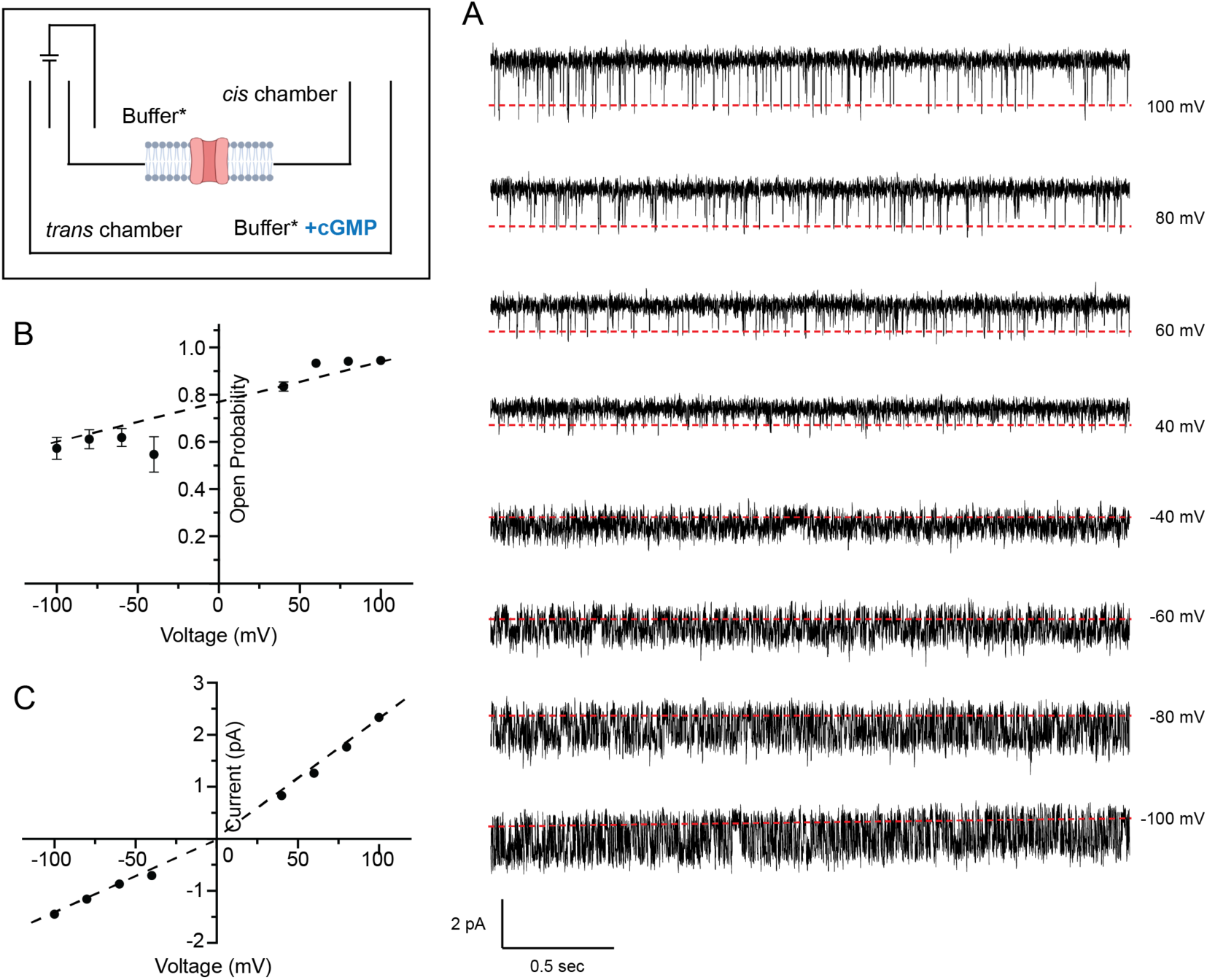
Characterization of single-channel activity of CNGA1 using planar lipid bilayer recordings. Left corner: Cartoon of a planar bilayer single-channel recording setup with two bilayer chambers (*cis* and *trans*) separated by a partition where a bilayer is painted (grey), shown as containing a channel (red). The *cis* chamber is where the liposomes are added, and the *trans* chamber is where the cGMP is added, to ensure recording only from channels oriented with the CNBD facing the *trans* chamber. (A) Representative single-channel traces recorded at voltages from –100 mV to +100 mV. The baseline is indicated by a red dotted line. All recordings were performed in the presence of 1 mM cGMP (saturating concentration). Scale bars: 2 pA, 0.5 s. (B) Voltage dependence of channel open probability extracted from data as in A. (C) Current–voltage (I–V) relationship of unitary currents. Bars show mean ± SEM (n=4). The trend lines in the plots are displayed as black dashed lines.

**Supplementary Figure 3.**
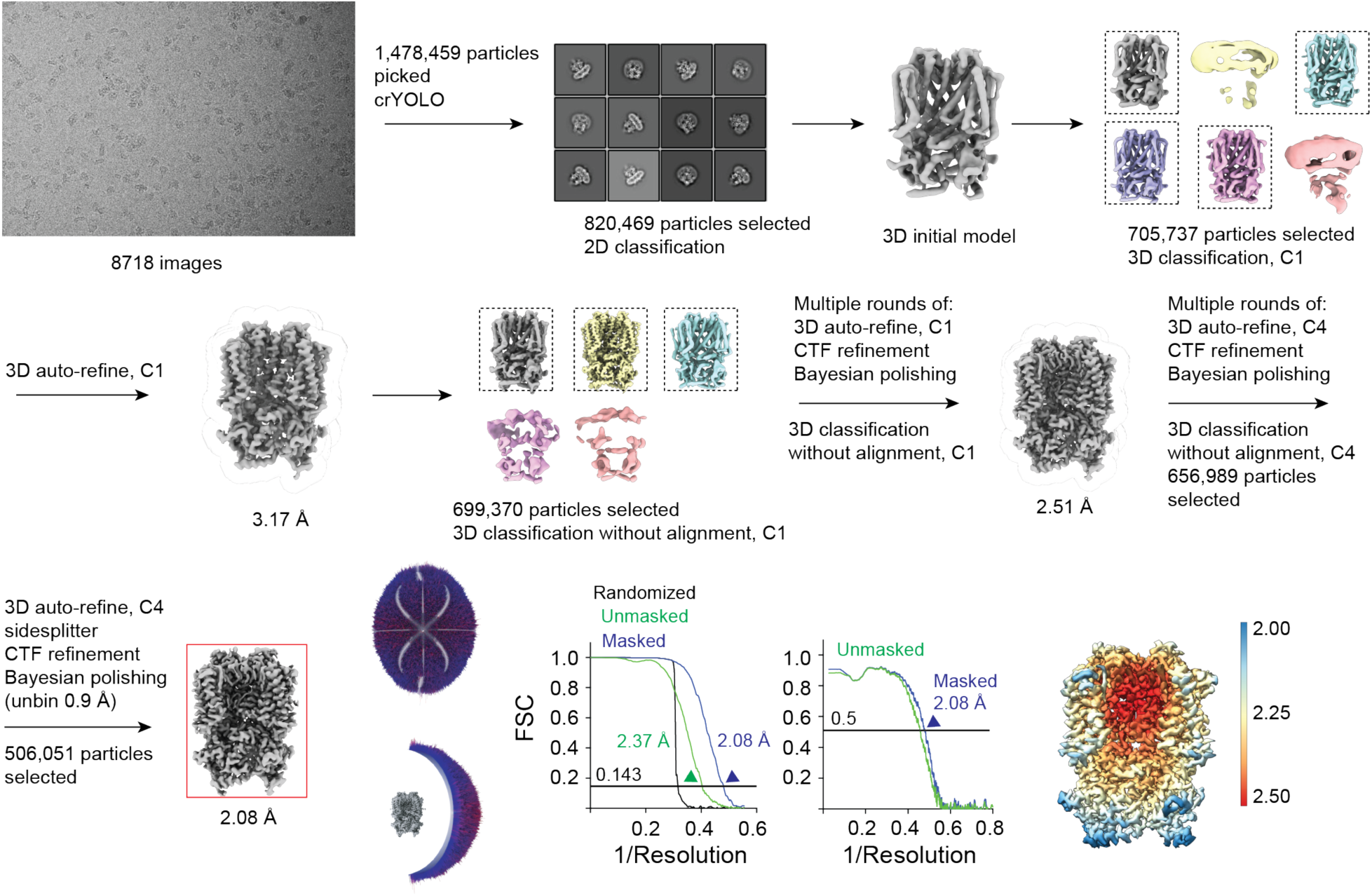
Cryo-EM data processing workflow for CNGA1 reconstituted into nanodiscs (65% DOPC, 11% POPE, 24% POPS) in the absence of cGMP. The workflow illustrates particle selection, 2D and 3D classification, and refinement. The final 3D reconstruction is highlighted with a red box.

**Supplementary Figure 4.**
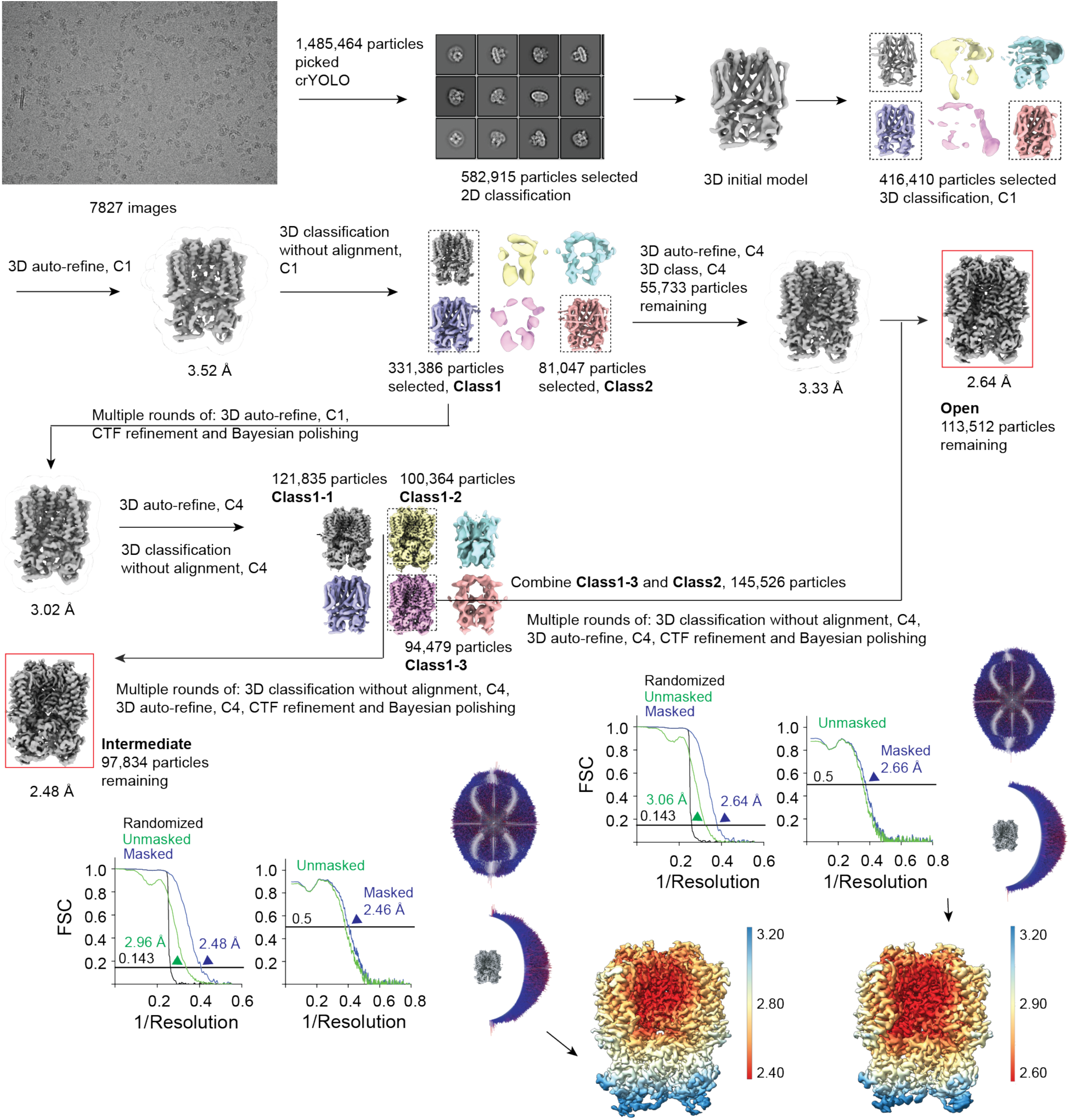
Cryo-EM data processing workflow for CNGA1 reconstituted into nanodiscs (65% DOPC, 11% POPE, 24% POPS) in the presence of 3 mM cGMP. The workflow illustrates particle selection, 2D and 3D classification, and refinement. The final 3D reconstructions, intermediate and open, are highlighted with red boxes.

**Supplementary Figure 5.**
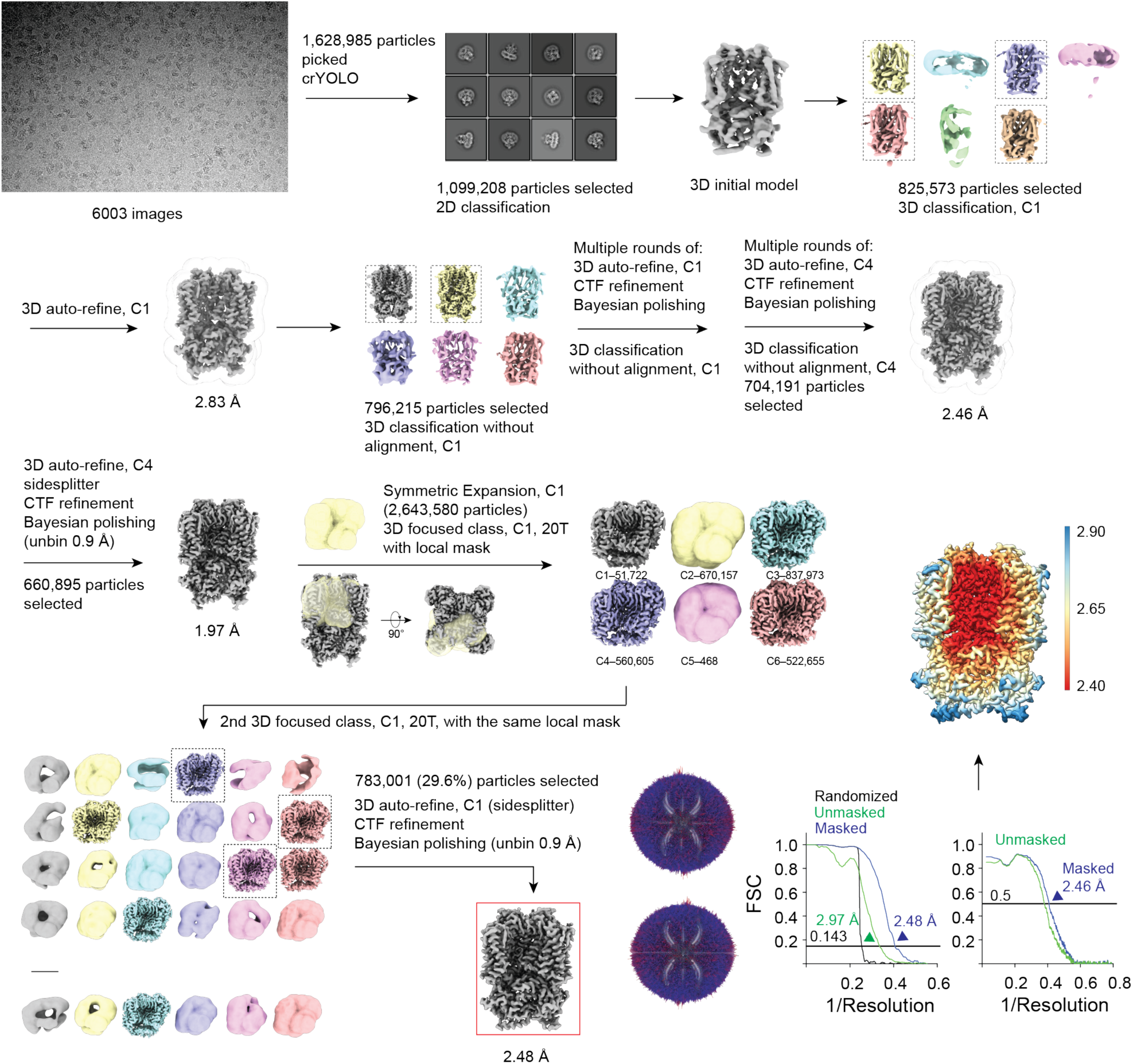
Cryo-EM data processing workflow for CNGA1 reconstituted into nanodiscs with 10 % brain PIP2 (55% DOPC, 11% POPE, 24% POPS, 10% brain PIP2) in the absence of cGMP. The workflow illustrates particle selection, 2D and 3D classification, refinement, symmetry expansion and focused classification using a local mask on one of the four lipid-binding pockets. The final 3D reconstruction is highlighted with a red box.

**Supplementary Figure 6.**
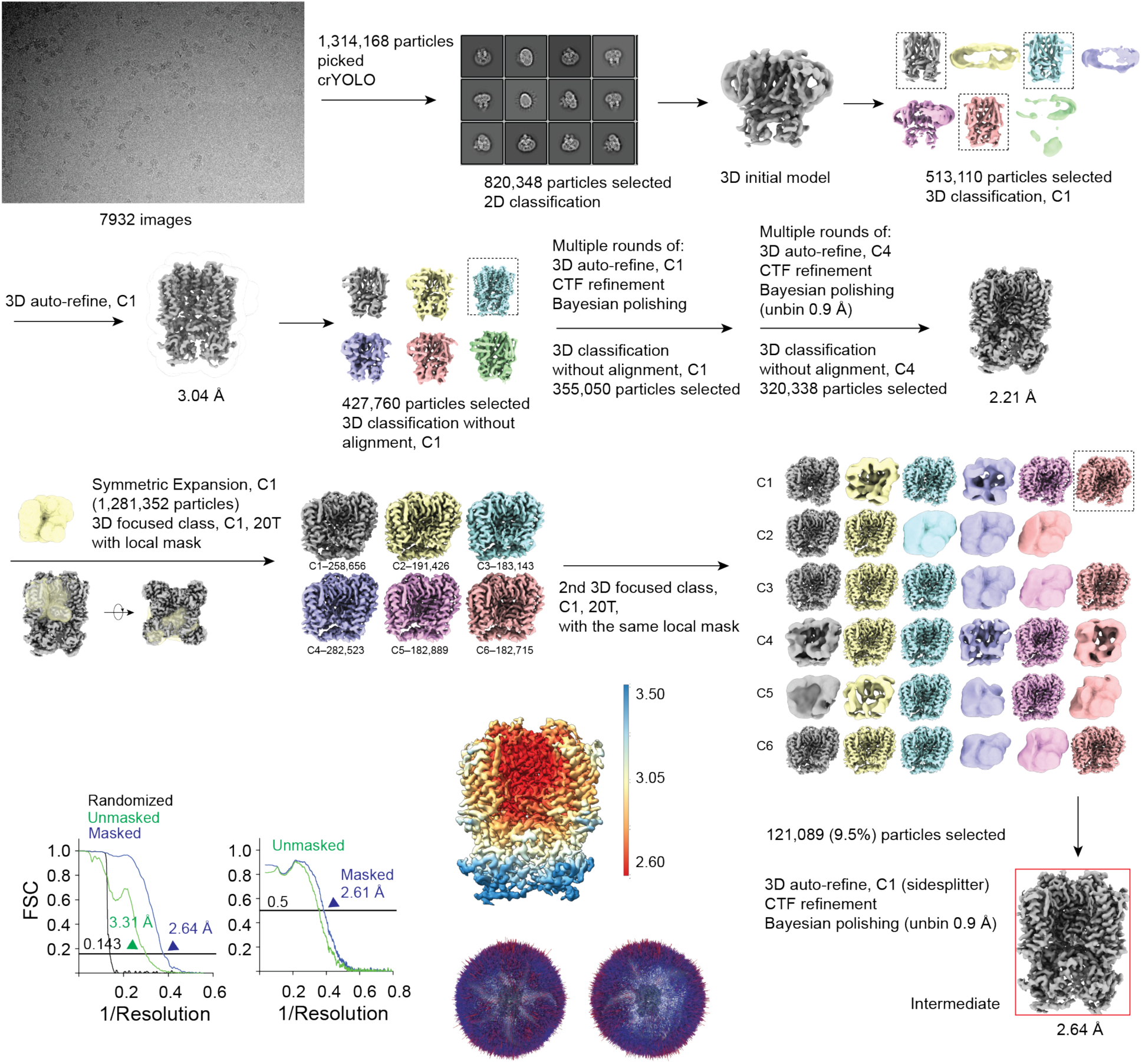
Cryo-EM data processing workflow for CNGA1 reconstituted into nanodiscs with 10% brain PIP2 (55% DOPC, 11% POPE, 24% POPS, 10% brain PIP2) in the presence of 3 mM cGMP. The workflow illustrates particle selection, 2D and 3D classification, refinement, symmetry expansion, and focused classification using a local mask on one of the four lipid-binding pockets. The final 3D reconstruction is highlighted with a red box.

**Supplementary Figure 7.**
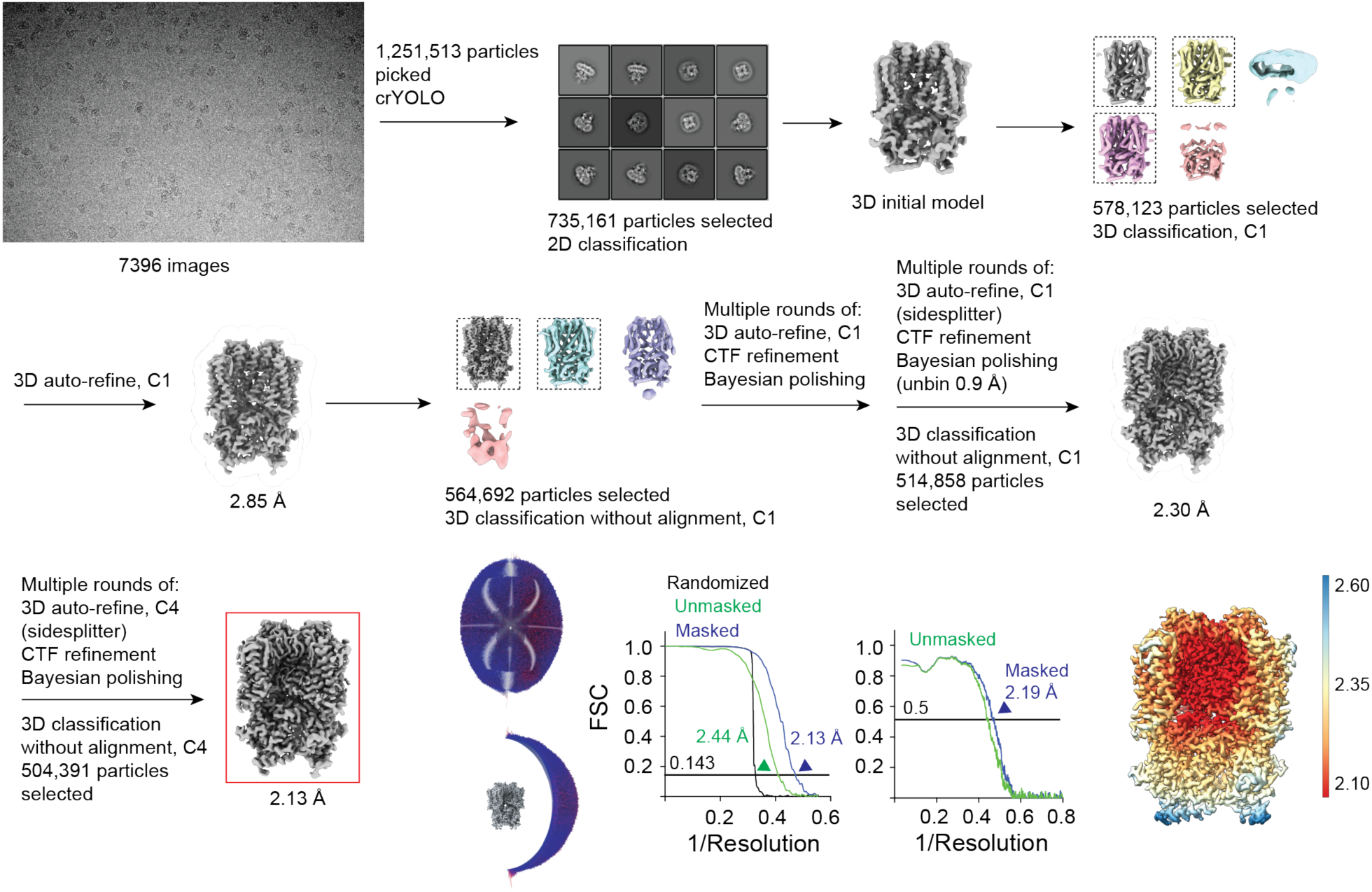
Cryo-EM data processing workflow for CNGA1 reconstituted into nanodiscs (65% DOPC, 11% POPE, 24% POPS) in presence of 3 mM diC8-PIP2 without cGMP. The workflow illustrates particle selection, 2D and 3D classification, and refinement. The final 3D reconstruction is highlighted with a red box.

**Supplementary Figure 8.**
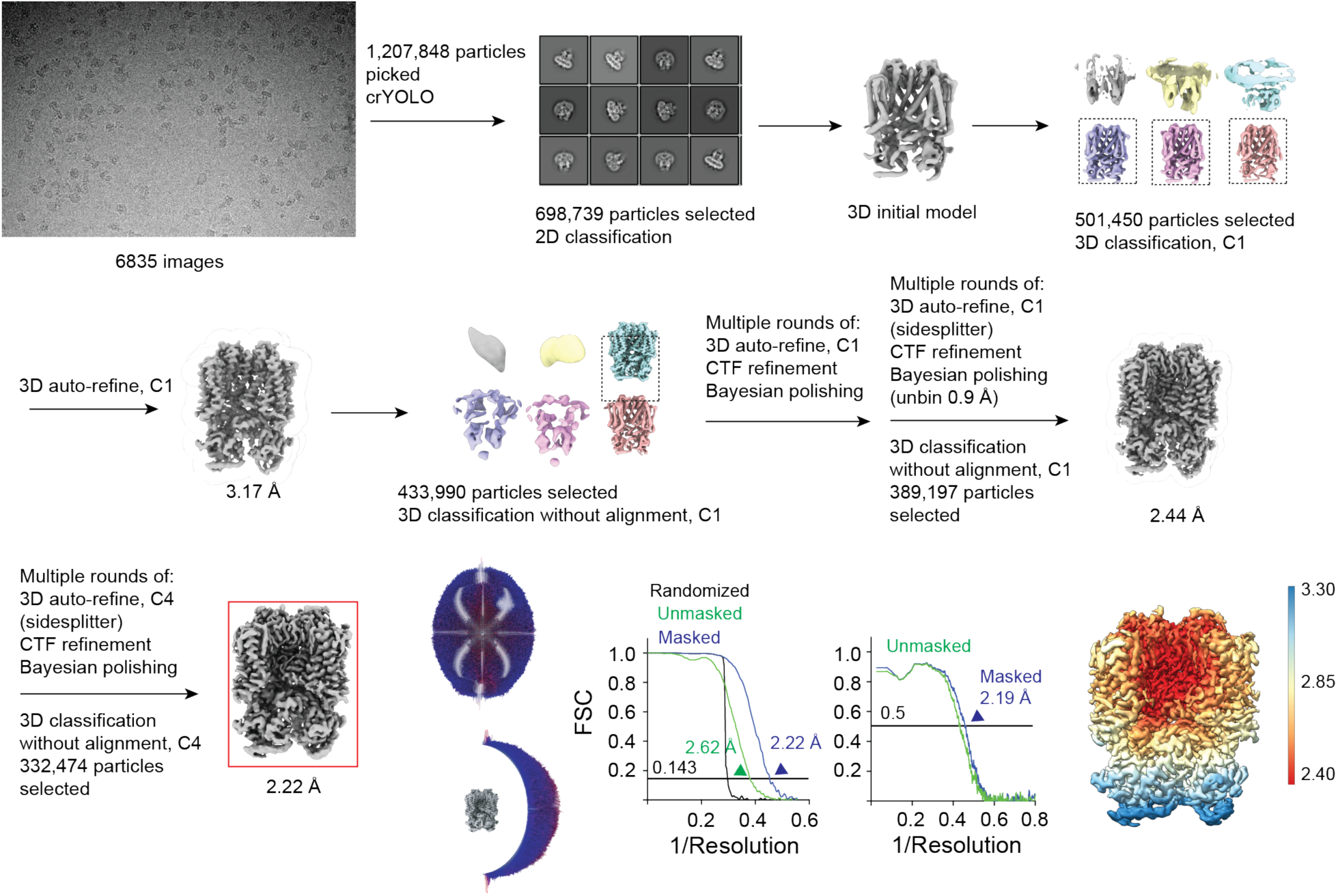
Cryo-EM data processing workflow for CNGA1 reconstituted into nanodiscs (65% DOPC, 11% POPE, 24% POPS) in presence of 3 mM diC8-PIP2 and 3 mM cGMP. The workflow illustrates particle selection, 2D and 3D classification, and refinement. The final 3D reconstruction is highlighted with a red box.

**Supplementary Figure 9.**
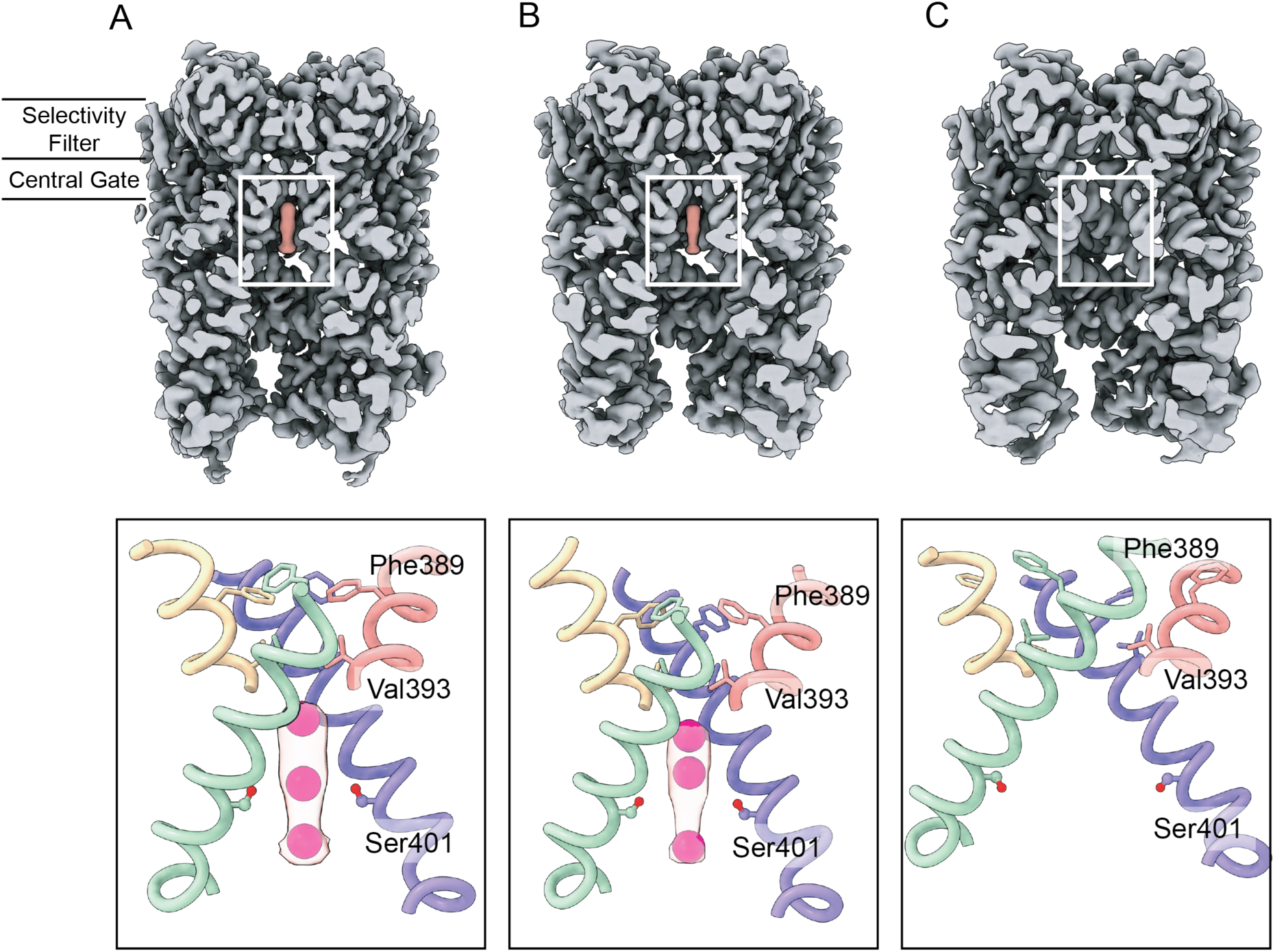
Newly observed densities below the central gate near Ser401. Panels show the closed (A), intermediate (B), and open (C) states of the channel, each displayed in overall, and side views. In the closed and intermediate states, additional densities (Potassium ions, magenta spheres) are observed immediately below the central gate, located near Ser401 and neighboring peptide backbones. In the open state, the gate is dilated, the distance from Ser401 to the central axis increases, and no corresponding densities are present. Each subunit is colored differently for clarity. In panel C, the additional densities present in the closed and intermediate structures are absent in the open conformation.

**Supplementary Figure 10.**
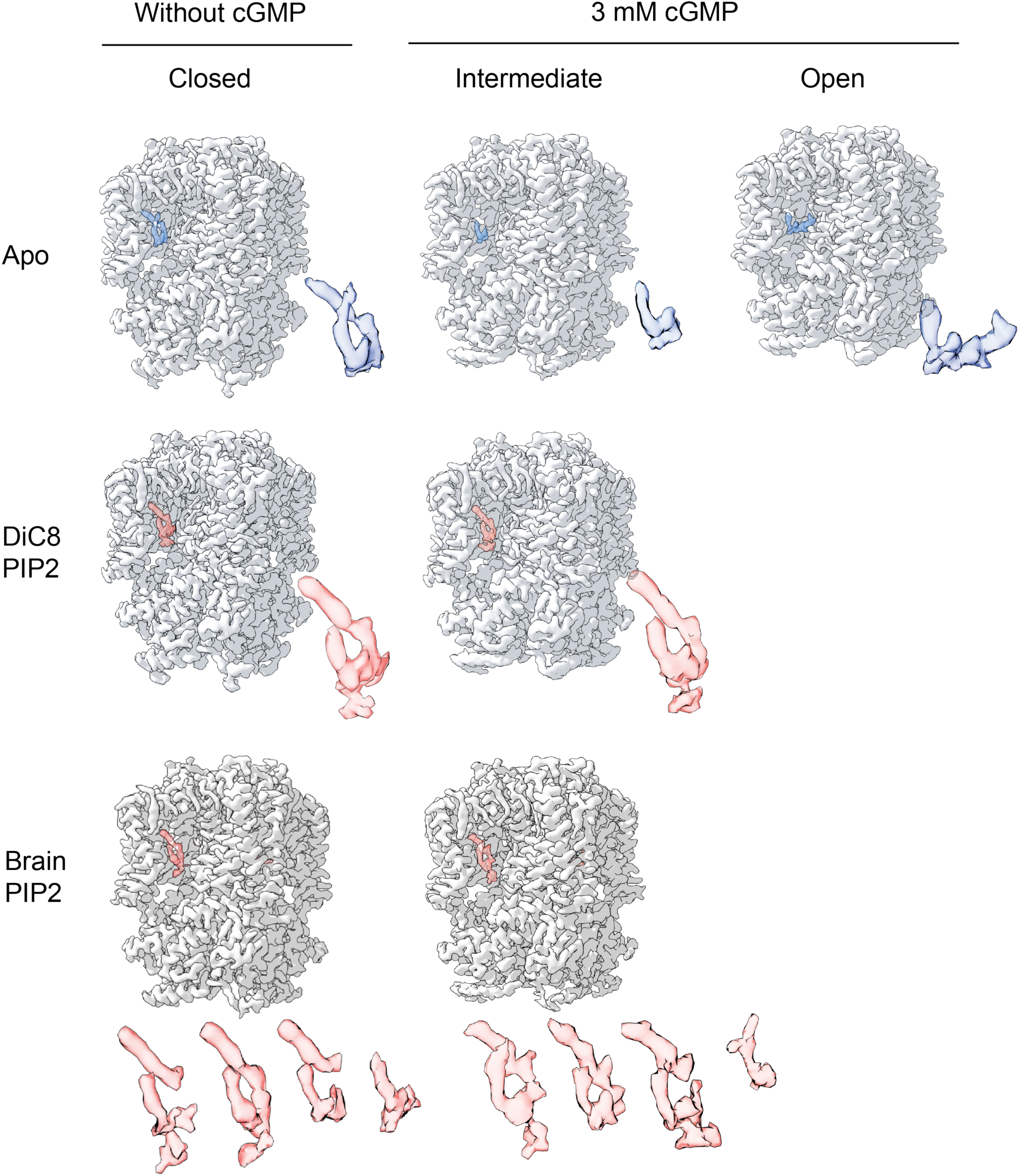
Comparison of lipid densities in the PIP2 pockets across our CNGA1 structures. PIP2 densities from CNGA1 structures determined in the presence of diC8-PIP2 or brain PIP2 are shown in red. Lipid-like densities observed in structures obtained without added PIP2 are shown in blue.

**Supplementary Figure 11.**
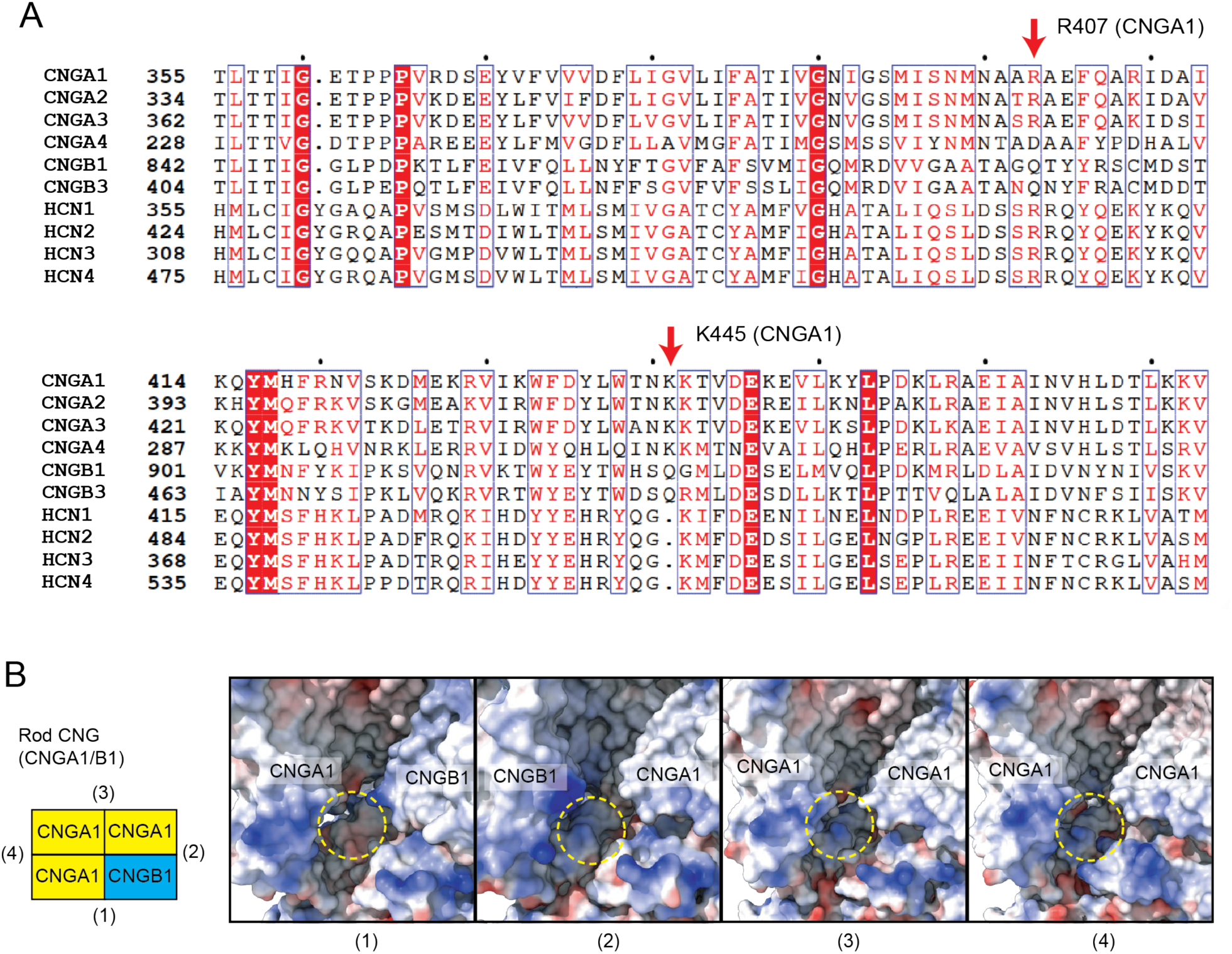
Sequence conservation across CNG/HCN channels and electrostatic features of rod CNG PIP2/lipid-binding sites. (A) Sequence alignment of CNG and HCN channel family members generated using Clustal Omega (https://www.ebi.ac.uk/Tools/msa/clustalo/) and visualized with ESPript 3.2 (https://espript.ibcp.fr/ESPript/ESPript/). Red arrows mark the residues corresponding to R407 and R445 in CNGA1. (B) Electrostatic surface representation of the four lipid-binding pockets in the rod CNGA1/CNGB1 heterotetramer structures (PDB ID: 7RH9). Each pocket is formed at the interface of two adjacent protomers, and the yellow dotted circles indicate the region where the PIP₂ headgroup would predominantly interact in the CNGA1 tetramer.

**Supplementary Table 1.**
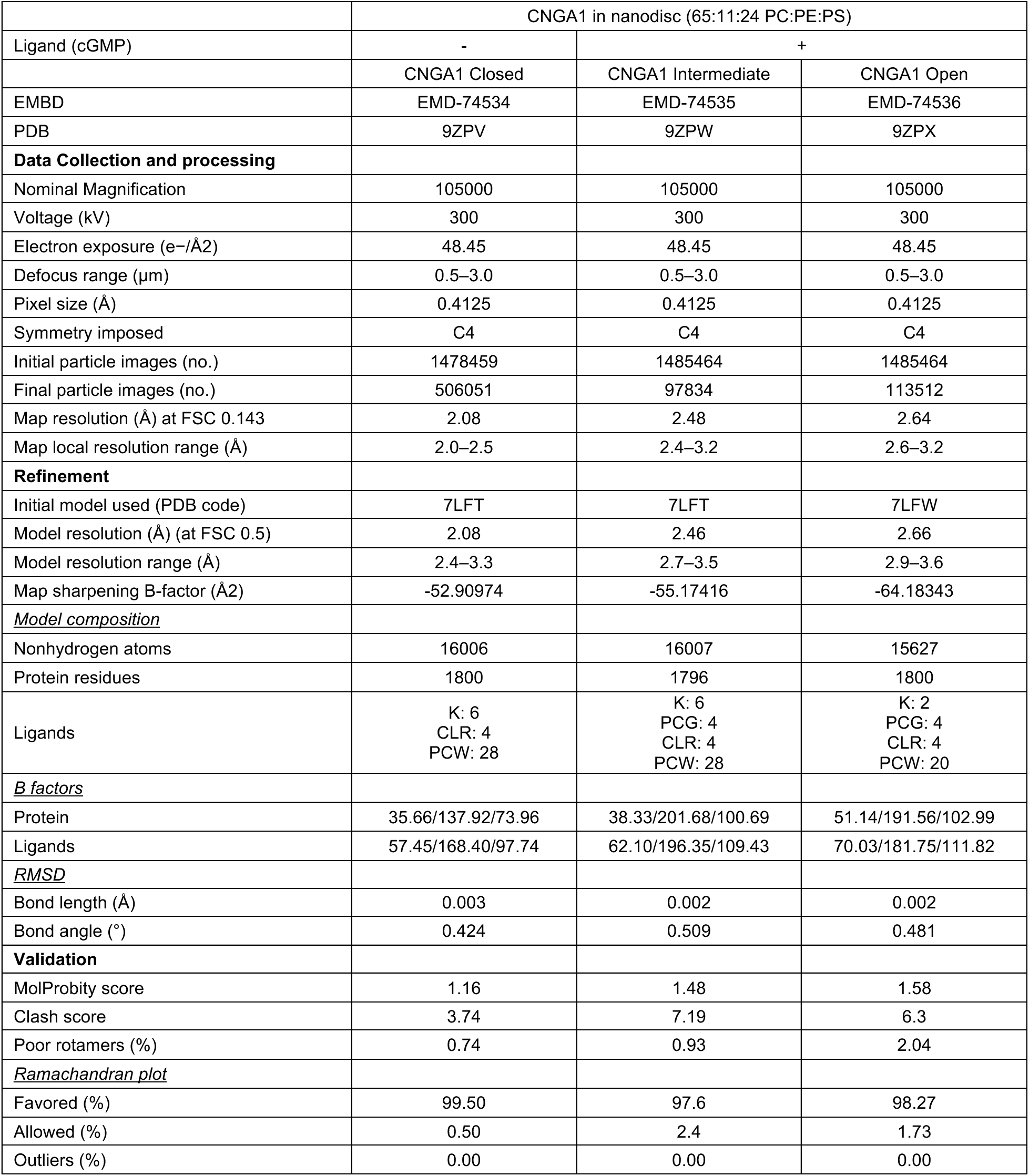
Cryo-EM data collection, refinement, and validation statistics for CNGA1 structures in nanodiscs.

|  | CNGA1 in nanodisc (65:11:24 PC:PE:PS) |  |  |
| --- | --- | --- | --- |
| Ligand (cGMP) | - | + |  |
|  | CNGA1 Closed | CNGA1 Intermediate | CNGA1 Open |
| EMBD | EMD-74534 | EMD-74535 | EMD-74536 |
| PDB | 9ZPV | 9ZPW | 9ZPX |
| **Data Collection and processing** |  |  |  |
| Nominal Magnification | 105000 | 105000 | 105000 |
| Voltage (kV) | 300 | 300 | 300 |
| Electron exposure (e-/Å<sup>2</sup>) | 48.45 | 48.45 | 48.45 |
| Defocus range (µm) | 0.5–3.0 | 0.5–3.0 | 0.5–3.0 |
| Pixel size (Å) | 0.4125 | 0.4125 | 0.4125 |
| Symmetry imposed | C4 | C4 | C4 |
| Initial particle images (no.) | 1478459 | 1485464 | 1485464 |
| Final particle images (no.) | 506051 | 97834 | 113512 |
| Map resolution (Å) at FSC 0.143 | 2.08 | 2.48 | 2.64 |
| Map local resolution range (Å) | 2.0–2.5 | 2.4–3.2 | 2.6–3.2 |
| **Refinement** |  |  |  |
| Initial model used (PDB code) | 7LFT | 7LFT | 7LFW |
| Model resolution (Å) (at FSC 0.5) | 2.08 | 2.46 | 2.66 |
| Model resolution range (Å) | 2.4–3.3 | 2.7–3.5 | 2.9–3.6 |
| Map sharpening B-factor (Å<sup>2</sup>) | -52.90974 | -55.17416 | -64.18343 |
| <u>Model composition</u> |  |  |  |
| Nonhydrogen atoms | 16006 | 16007 | 15627 |
| Protein residues | 1800 | 1796 | 1800 |
| Ligands | K: 6<br>CLR: 4<br>PCW: 28 | K: 6<br>PCG: 4<br>CLR: 4<br>PCW: 28 | K: 2<br>PCG: 4<br>CLR: 4<br>PCW: 20 |
| <u>B factors</u> |  |  |  |
| Protein | 35.66/137.92/73.96 | 38.33/201.68/100.69 | 51.14/191.56/102.99 |
| Ligands | 57.45/168.40/97.74 | 62.10/196.35/109.43 | 70.03/181.75/111.82 |
| <u>RMSD</u> |  |  |  |
| Bond length (Å) | 0.003 | 0.002 | 0.002 |
| Bond angle (°) | 0.424 | 0.509 | 0.481 |
| **Validation** |  |  |  |
| MolProbity score | 1.16 | 1.48 | 1.58 |
| Clash score | 3.74 | 7.19 | 6.3 |
| Poor rotamers (%) | 0.74 | 0.93 | 2.04 |
| <u>Ramachandran plot</u> |  |  |  |
| Favored (%) | 99.50 | 97.6 | 98.27 |
| Allowed (%) | 0.50 | 2.4 | 1.73 |
| Outliers (%) | 0.00 | 0.00 | 0.00 |

**Supplementary Table 2.** Cryo-EM data collection, refinement and validation statistics for CNGA1 structures in nanodiscs with brain PIP2 and with diC8-PIP2.

|  | CNGA1 in nanodisc with Brain PIP2<br>(55:11:24:10 PC:PE:PS:Brain PIP2) |  | CNGA1 in nanodisc (65:11:24 PC:PE:PS)<br>with diC8 PIP2 |  |
| --- | --- | --- | --- | --- |
| Ligand (cGMP) | - | + | - | + |
|  | CNGA1 Closed | CNGA1<br>Intermediate | CNGA1 Closed | CNGA1<br>Intermediate |
| EMBD | EMD-74537 | EMD-74538 | EMD-74539 | EMD-74540 |
| PDB | 9ZPY | 9ZPZ | 9ZQ0 | 9ZQ1 |
| **Data Collection and processing** |  |  |  |  |
| Nominal Magnification | 105000 | 105000 | 105000 | 105000 |
| Voltage (kV) | 300 | 300 | 300 | 300 |
| Electron exposure (e-/Å<sup>2</sup>) | 48.56 | 48.56 | 48.48 | 48.48 |
| Defocus range (µm) | 0.6–2.5 | 0.6–2.5 | 0.6–2.4 | 0.6–2.4 |
| Pixel size (Å) | 0.4125 | 0.4125 | 0.4125 | 0.4125 |
| Symmetry imposed | C1 | C1 | C4 | C4 |
| Initial particle images (no.) | 1628985 | 1314168 | 1251513 | 1207848 |
| Final particle images (no.) | 195750 | 30272 | 504391 | 332474 |
| Map resolution (Å) at FSC 0.143 | 2.48 | 2.64 | 2.13 | 2.22 |
| Map local resolution range (Å) | 2.4–2.9 | 2.6–3.5 | 2.1–2.6 | 2.4–3.3 |
| **Refinement** |  |  |  |  |
| Initial model used (PDB code) | 7LFT | 7LFT | 7LFT | 7LFT |
| Model resolution (Å) (at FSC 0.5) | 2.45 | 2.61 | 2.11 | 2.19 |
| Model resolution range (Å) | 2.6–3.5 | 2.7–3.4 | 2.5–3.0 | 2.6–3.4 |
| Map sharpening B-factor (Å<sup>2</sup>) | -48.15762 | -42.63815 | -53.98723 | -52.20838 |
| <u>Model composition</u> |  |  |  |  |
| Nonhydrogen atoms | 16048 | 16102 | 16138 | 16192 |
| Protein residues | 1800 | 1796 | 1800 | 1796 |
| Ligands | K: 5<br>PT5: 2<br>CLR: 4<br>PCW: 28 | K: 5<br>PCG: 2<br>PT5: 4<br>CLR: 4<br>PCW: 28 | K: 5<br>PIO: 4<br>CLR: 4<br>PCW: 28 | K: 5<br>PCG: 4<br>PIO: 4<br>CLR: 4<br>PCW: 28 |
| <u>B factors</u> |  |  |  |  |
| Protein | 35.47/157.48/76.51 | 30.19/193.25/94.54 | 27.69/137.08/66.99 | 34.99/202.98/97.28 |
| Ligands | 55.08/205.10/104.96 | 50.19/199.43/98.99 | 51.20/220.52/99.20 | 58.46/226.12/109.43 |
| <u>RMSD</u> |  |  |  |  |
| Bond length (Å) | 0.002 | 0.002 | 0.002 | 0.002 |
| Bond angle (°) | 0.411 | 0.472 | 0.424 | 0.492 |
| **Validation** |  |  |  |  |
| MolProbity score | 1.40 | 1.72 | 1.22 | 1.61 |
| Clash score | 5.11 | 7.55 | 4.47 | 7.27 |
| Poor rotamers (%) | 1.48 | 1.60 | 0.80 | 1.30 |
| <u>Ramachandran plot</u> |  |  |  |  |
| Favored (%) | 98.21 | 97.26 | 98.83 | 97.37 |
| Allowed (%) | 1.79 | 2.74 | 1.17 | 2.63 |
| Outliers (%) | 0.00 | 0.06 | 0.00 | 0.00 |

## Notes

### Competing Interest Statement

The authors have declared no competing interest.

